# Localization and connectivity of rodent equivalent of the primate posterior cingulate cortex (area 23)

**DOI:** 10.1101/2023.01.11.523665

**Authors:** Xiao-Jun Xiang, Sheng-Qiang Chen, Xue-Qin Zhang, Chang-Hui Chen, Shun-Yu Zhang, Hui-Ru Cai, Song-Lin Ding

## Abstract

The posterior cingulate cortex (mainly area 23) in human and non-human primates is a critical component of the default mode network and is involved in many neurological and neuropsychiatric diseases such as Alzheimer’s disease, autism, depression, attention deficit hyperactivity disorder and schizophrenia. However, cingulate area 23 has not yet identified in rodents and other lower mammals and this makes modeling related circuits and diseases in rodents very difficult. Using a comparative approach and unique connectional patterns the present study has uncovered the location and extent of rodent equivalent of the primate cingulate area 23. Like in monkeys, area 23 but not adjoining retrosplenial and visual areas in the rats and mice displays strong reciprocal connections with the anteromedial thalamic nucleus. Rodent area 23 also reciprocally connects with the medial pulvinar and claustrum as well as with the anterior cingulate, granular retrosplenial, medial orbitofrontal, postrhinal, and visual and auditory association cortices. The rodent A23 also projects to the subcortical effectors such as the dorsal striatum, ventral lateral geniculate nucleus, zona incerta, pretectal nucleus, superior colliculus, periaqueductal gray, and brainstem reticular formation. All these connectional findings support the versatility of area 23 in the integration and modulation of multimodal information underlying spatial processing, episodic memory, self-reflection, attention, value assessment and many adaptive behaviors. Additionally, this study also suggests that the rodents can be used to model primate and human area 23 in future structural, functional, pathological and neuromodulation studies.

## 1. Introduction

The posterior cingulate gyrus in human and non-human primates (NHP) mainly contains the retrosplenial cortex (RS), which usually includes Brodmann’s areas 29 and 30 (A29 and A30, respectively), and the posterior cingulate cortex (PCC), which contains a large area 23 (A23) and a smaller area 31 (A31) (Brodmann, 1909; Vogt et al., 1987; Morecraft et al., 2004; Ding et al., 2016). Topographically, A30, A23 and A31 are located dorsal to A29, A30 and A23, respectively, and extend together as an arch around the splenium of the corpus callosum (CCS) (Vogt et al., 1987; Morris et al., 1999; Morecraft et al., 2004; Vogt et al., 2005). As in human and NHP, rodent A29 [i.e., granular RS (RSg)] is similarly located dorsal and caudal to the CCS while A30 [i.e., agranular RS (RSag)] is located dorsolateral to A29 (Sugar et al., 2011; Vogt and Paxinos 2014). However, homologous PCC has not been reported in rodent literature so far. Instead, the region lateral to A30 (RSag) has been treated as a lateral agranular region of the RS (RSagl, e.g., Swanson, 2004; Hu et al., 2020; Wang et al., 2020) or a part of the medial secondary visual cortex (area 18b or V2MM, e.g., Vogt and Miller, 1983; Sripanidkulchai and Wyss, 1986; van Groen et al., 1999; Paxinos and Franklin, 2012). The reasons for the difficulty of identifying the rodent equivalent of the PCC may include both historical and cytoarchitectonic aspects. Historically, the PCC has not been identified in lower mammals since Brodmann’s comparative mapping of the cerebral cortex (Brodmann, 1909; Vogt and Paxinos, 2014). Cytoarchitectonically, the existence (for the PCC) or lack (for the RS) of an inner granular layer 4 was used as the main criteria to define the PCC and RS, respectively (Brodmann, 1909; Morris et al., 1999; Vogt et al., 2005). Accordingly, the region located immediately lateral to A30 in rodents was not treated as the PCC since the region did not appear to have a granular layer 4 (Vogt and Paxinos 2014).

However, in modern literature, brain structures/regions have also been defined using other important organizational features such as connectional patterns (Fudge et al., 2004; Gehrlach et al, 2020) and molecular signatures (Wallen-Mackenzie et al., 2020; Chen et al., 2022; Ding et al., 2022) as well as using combination of multimodal data (Ding and van Hoesen, 2010, 2015; Ding, 2022). In this study, we aim to investigate the region located immediately lateral to A30 in rat and mouse brains to determine whether this region possess similar connectional patterns as the monkey PCC (mainly A23) does. A23 but not adjoining A30 in the monkey brains was reported to receive strong inputs from the anteromedial thalamic nucleus (AM; Baleydier and Mauguiere, 1980; Vogt et al. 1987; Morris et al., 1999). Based on this finding we speculate that if the RSag (A30) and the RSagl (or part of it) in rodents receives, respectively, few or strong AM inputs, the latter region is probably the rodent equivalent of the monkey and human A23. Furthermore, if the rodent equivalent of A23 (termed A23~ in this study) identified based on the AM inputs is indeed the homolog of primate A23, A23~ should has similar brain-wide connectivity to that of the monkey A23 and should also be distinguishable from adjoining A30.

The results of the present study have confirmed above speculations and thus uncovered the long-lasting mystery about whether rodent homolog of primate A23 exists. The connectivity of A23~ revealed in the present study would provide important insights into both evolutional and functional roles of A23~. Moreover, the identification of rodent A23~ also suggests that rodents can be used as animal models for future studies of the NHP and human PCC (mainly A23). In human, dysfunctions of the PCC have been observed in many neurological and mental diseases such as Alzheimer’s disease, traumatic brain injury, autism, depression, attention deficit hyperactivity disorder, addiction, and schizophrenia (Bluhm et al., 2009; Leech and sharp, 2014; Rolls, 2019; Zhang and Volkow, 2019). Detailed studies are needed in rodent models to gain more insights into the neural circuits and mechanisms of these diseases and the development of interference strategies.

## 2. Materials and Methods

### Animals

Forty adult Sprague-Dawley rats of both sexes weighing 280–310 g (Beijing Vital River Laboratory Animal Technology Co., Ltd., Beijing, China) were used in this study. All the animals were placed in the same environment with suitable temperature and controlled light, as well as free access to food and water. All surgery operations were performed under the deep anesthesia to alleviate their pain. All experimental procedures were followed in accordance with the protocols that have been approved by the Institutional Animal Care and Use Committee.

### Surgery Procedure and Tracer Injections

The surgeries were carried out when the rats were deeply anesthetized with sodium pentobarbital (40 mg/kg, i. p.). The rat was fixed in a stereotaxic frame and the head hairs were shaved and a midline incision was made after skin disinfection to expose the surgical field. After the bregma and lambda were adjusted to the same horizontal level, two bone windows (one on each side) were drilled on the skull over the target regions following the coordinates derived from the rat brain atlas of (Paxinos and Watson, 2013). Then 0.1 μl of 10% biotinylated dextran amine (BDA, 10,000 MW, ThermoFisher Scientific, Waltham, MA, United States) or 4% FluroGold (FG, Fluorochrome Inc., Denver, CO, United States) was pressure injected into the target brain regions of one hemisphere using a 0.5-μl Hamilton syringe. The needle was hold for 10min after the tracer injection and then pulled out slowly. Finally, the incisions were sutured, and the rats were returned to their home cages after recovery on the warm bed.

### Brain Preparation

7-10 days after the surgeries, the rats were deep anesthetized and perfused with 0.9% saline and 4% paraformaldehyde (PFA) in chilled 0.05M phosphate buffer (PB, pH 7.3) in sequence. Then the brains were removed and immersed in the 4% PFA at 4°**C** overnight, and then cryoprotected in the PB containing 15 and 30% sucrose in succession for 3-4 days until the brains sank to the bottom. Each brain was separated into two halves with a midline cut. And each hemisphere was cut into 40μm-thick coronal sections using a freezing microtome.

### Histochemistry for BDA Tracing

The procedure for BDA histochemistry has been described in our previous study (Lu et al., 2020). In brief, the sections derived from the hemisphere with BDA injections were thoroughly rinsed (at least three times 10 min each) in 0.05M PB. Then the sections were incubated in 0.3% Triton X-100 in 0.05M PB for 1h and in Streptavidin-Biotin Complex solution (SABC kit, Boster Biological Technology) for 3h at room temperature in sequence. After rinse in 0.05M PB for three times, the sections were visualized with 0.05M PB containing 0.05% 3, 3-diaminobenzidine (DAB). Then the sections were mounted on glass slides, dehydrated in gradient alcohol and xylene, and finally coverslipped.

### Immunohistochemistry for FG Tracing

Sections from the cases with FG injections were observed under an epifluorescent microscope (Leica DM6B) or stained with immunohistochemistry (IHC) following the guide of standard procedures. For the IHC, the sections were rinsed in 0.05M PB for three times (5 min each) and incubated with 3% hydrogen peroxide for 10 min in room temperature. After blocking in 5% BSA for 60 min, the sections were incubated with 0.05M PB containing 0.3% triton X-100 and the primary antibody (rabbit anti-FG, AB153-I, 1:10000, Sigma-Aldrich) at 4°Covernight. After thorough rinses, the sections were incubated with the secondary antibody solution (biotinylated goat anti-mouse/rabbit IgG, Boster Biological Technology) and then the Streptavidin-Biotin Complex solution (SABC kit, Boster Biological Technology) for 60 min each. The sections were visualized in PB containing 0.05% DAB. Finally, the sections were mounted on chrome alum and gelatin-coated slides, dehydrated in gradient alcohol and xylene, and coverslipped.

### Image Acquisition and Processing

The images of stained sections were acquired with a scanner (Aperio CS2, Leica). All the selected images were processed with Adobe photoshop for image cropping, brightness and contrast adjustment, image composing, and structural annotation.

### Cell counts of the labeled neurons in the anterior thalamic nuclei

To compare the numbers of FG or BDA retrogradely labeled neurons in the AD, AV and AM after A23~ and A30 injections in the rats, the numbers of these neurons in the three structures were counted on sequential sections containing these structures. These sections were grouped into five A-P levels and each level includes two closely adjacent sections. Cell counts were performed using Image J and the statistics were done using one-way ANOVA and Tukey test (for each of the two pairs) or two-tailed unpaired t-test (for comparing cell numbers in the AM, AD or AV between A30 and A23 injections).

## 3. Results

### 3.1 Cytoarchitecture and topography of A23 in primates and rodents

Human A23 has similar cytoarchitecture and topography to monkey A23 (e.g., Vogt et al., 2001). In monkey brains, A23 is typically located dorsal to A30 while A29 adjoins A30 ventrally (e.g., Fig. 1A, B). A29 is characterized by densely packed small granular cells in its superficial layers 2 and 3 (i.e., outer granular cells), which are not distinguishable from each other (outlined in Fig. 1C). The thick deep layers (layers 5-6) of A29 are mainly composed of larger pyramidal cells (Fig. 1C). It is noted that a layer of larger cells is observed superficial to the small-celled layers 2-3 of A29 and is termed A30e (30e in Fig. 1C; e for extension) since A30e appears to be a lateral extension of the superficial layers of A30 compared to rodent A29 (Fig. 1D-F). Both A30 and A23 of the monkey brains do not contain the densely packed outer granular cells, but A23 has a thin but clear inner granular layer (layer 4) with no or faint *Enc1* expression. Both A30 and A23 have a thick layer 5 and an even thicker layer 6 with strong *Enc1* expression (Fig. 1A, B).

**Figure 1.**
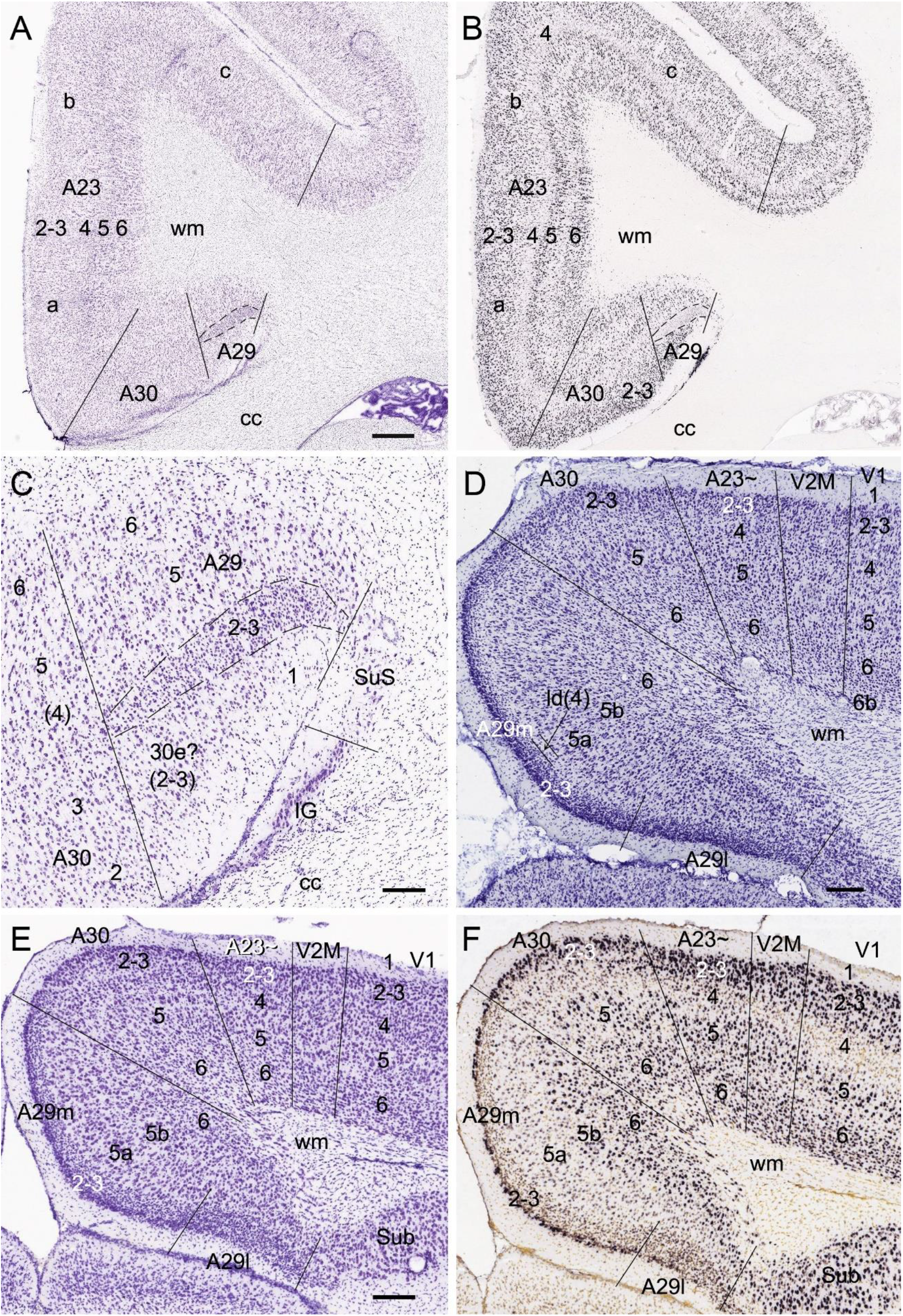
Cytoarchitecture and topography of area 23 in the monkey and rodent. For all panels, Arabic numbers (1–6) indicate cortical layers while solid straight lines mark the regional borders. For orientation of each section, medial is at the left and dorsal is at the top. (A, B) Locations, layers, subdivisions, and topography of areas 29, 30 and 23 (A29, A30 and A23) in the monkey. Two closely adjacent sections were stained for Nissl substances (A) and *Enc1* expression (ISH; B). A30 is typically located ventrally to A23 and dorsally to A29. Note that no or faint *Enc1* expression is detected in the thin but clear layer 4 (inner granular layer) whereas strong *Enc1* expression is seen in the thick layers 5 and 6 of A23. (C) Higher magnification view of A29 and A30 from panel (A). Layers 2-3 of A29 are outlined by dashed lines and are densely packed with small granular cells (outer granular layer). Note that a layer of larger cells, which is probably a lateral extension of the superficial layers of A30 (e for extension), is observed superior to layers 2-3 of A29. (D) Locations, layers, subdivisions, and topography of A29, A30 and A23 in the rat. The arrow indicates a cell-sparse zone (ld, lamina dissecans), which separates layers 2–3 and 5. (E, F) Locations, layers, subdivisions, and topography of A29, A30 and A23 in the mouse, showing on two closely adjacent sections stained for Nissl substances (E) and *Enc1*-ISH (F). Note that in the rodents A30 is typically located medially to A23 and dorsally to A29, and layers 2 and 3 of A29 is distinguishable from each other. Note that no extra layer appears superficial to layers 2-3 of A29 in the rodents. Also note the existence of a thick extra sublayer of layer 5 (layer 5a) in the mouse A29 with faint *Enc1* expression (F) although strong *Enc1* expression is observed in typical layer 5 of A29, A30 and A23 (layer 5b; F) as in the monkey (B). Both Layers 5a and 5b of A29 in the mouse express pan-layer 5 marker genes such as *Etv1*, *Rbp4* and *Adcyap1* (see Hu et al., 2020). Bars: 790 μm in (A) for (A, B); 197 μm in (C); 184 μm in (D); 210 μm in (E) for (E, F).

Like in the monkeys, layers 2-3 of A29 in the rat (Fig. 1D) and mouse (Fig. 1E) also consist of densely packed outer granular cells. However, layers 2 and 3 of A29 in rodents are distinguishable from each other. Layer 3 is densely packed with small cells and negative for *Enc1* while layer 2 is darkly stained and even more densely packed and is positive for *Enc1* (Fig. 1D-F). Unlike in the monkeys, no extra A30e appears superficial to layers 2-3 of A29 in rodents (Fig. 1D-F). Interestingly, A29 appears to have an extra sublayer of layer 5 (layer 5a) in rodents, which is thick and negative for *Enc1* compared to typical layer 5 (layer 5b), which is continuous with layer 5 in A30, A23 and neocortex and positive for *Enc1* (Fig. 1D-F). Both Layers 5a and 5b of A29 in the mice express pan-layer 5 marker genes such as *Etv1*, *Rbp4* and *Adcyap1* (see Hu et al., 2020). A29 in the rodents displays a thinner layer 6 compared to the monkeys. Topographically, A30 in the rodents is located dorsolateral to A29 while A23~ adjoins A30 laterally (Fig. 1D, E). A30 in the rodents has less densely packed layers 2-3, which are *Enc1* positive and not distinguishable from each other, and thick layers 5 and 6 with positive *Enc1* (Fig. 1D-F). A23~ in the rodents has slightly thicker layers 2-3 and slightly thinner layers 5-6 compared to A30 as well as larger cells in layer 6 compared to adjoining V2M. It is obvious on coronal sections that A23 in the primates is much wider/larger in size than A29 (e.g., Fig. 1A) while A23~ in the rodents is much narrower/smaller in size than A29 although the relative size of A30 appears comparable in both primates and rodents. In general, it is difficult to differentiate A30 from A23~ in the rodents based on Nissl staining.

### 3.2 A23~ but not A30 receives major inputs from anteromedial thalamic nucleus in rats

As mentioned above, the monkey AM projects to A23 while the AV and AD projects to A29 and A30 (Baleydier and Mauguiere, 1980; Vogt et al., 1987; Morris et al., 1999). To determine whether A23~ receives major inputs from the AM in the rats we performed FG injections in the region corresponding to A23~ described above. After successful FG injections (4 cases), many retrogradely labeled neurons were found in the AM with no or few in AD and AV. For example, following one single FG injection in A23~ (Fig. 2A) the labeled neurons are distributed across the anterior-posterior (A-P) extent of the AM with no or few in other thalamic regions including the AD, AV and PT (Fig. 2B-H). Some labeled cells were also observed in the ventromedial thalamic nucleus (VM) underneath the posterior AM.

**Figure 2.**
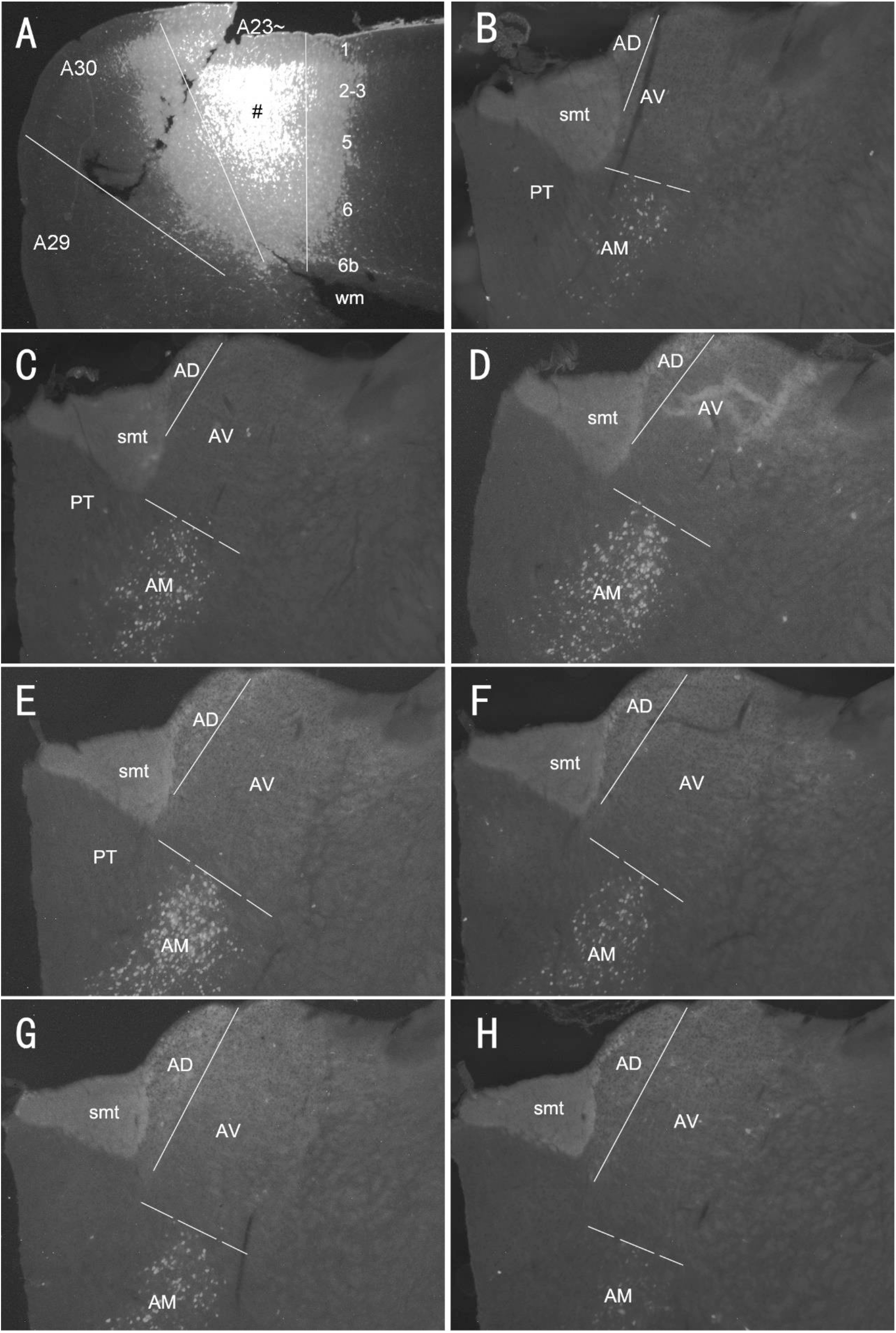
Labeled neurons in the AM following FG injection in the rat A23~. (A). One FG injection site (#) in A23~. (B-H) Sequential coronal sections from the anterior (B) to posterior (H) levels showing FG labeled neurons in the AM, AD and AV. The borders between the AD and AV and between AV and AM are indicated by a solid line and a dashed line, respectively. Note that most of the labeled neurons are concentrated in the AM across the A-P extent with no or few in the AD and AV. Bar: 500 μm in (A) for all panels.

In contrast, following FG or BDA injections in A30 (6 cases), retrogradely labeled neurons were observed mostly in AD and AV with no or few in the AM. For example, when the FG injections were placed in A30 (4 cases with 2 cases slightly involved in adjoining A29), FG labeled neurons are mostly seen in the AD and AV with no or few labeled neurons in the AM (e.g., Fig. 3A-H). These results indicate that both the AD and AV but not the AM project to A30. These findings are consistent with those in the monkeys (e.g., Vogt et al., 1987).

**Figure 3.**
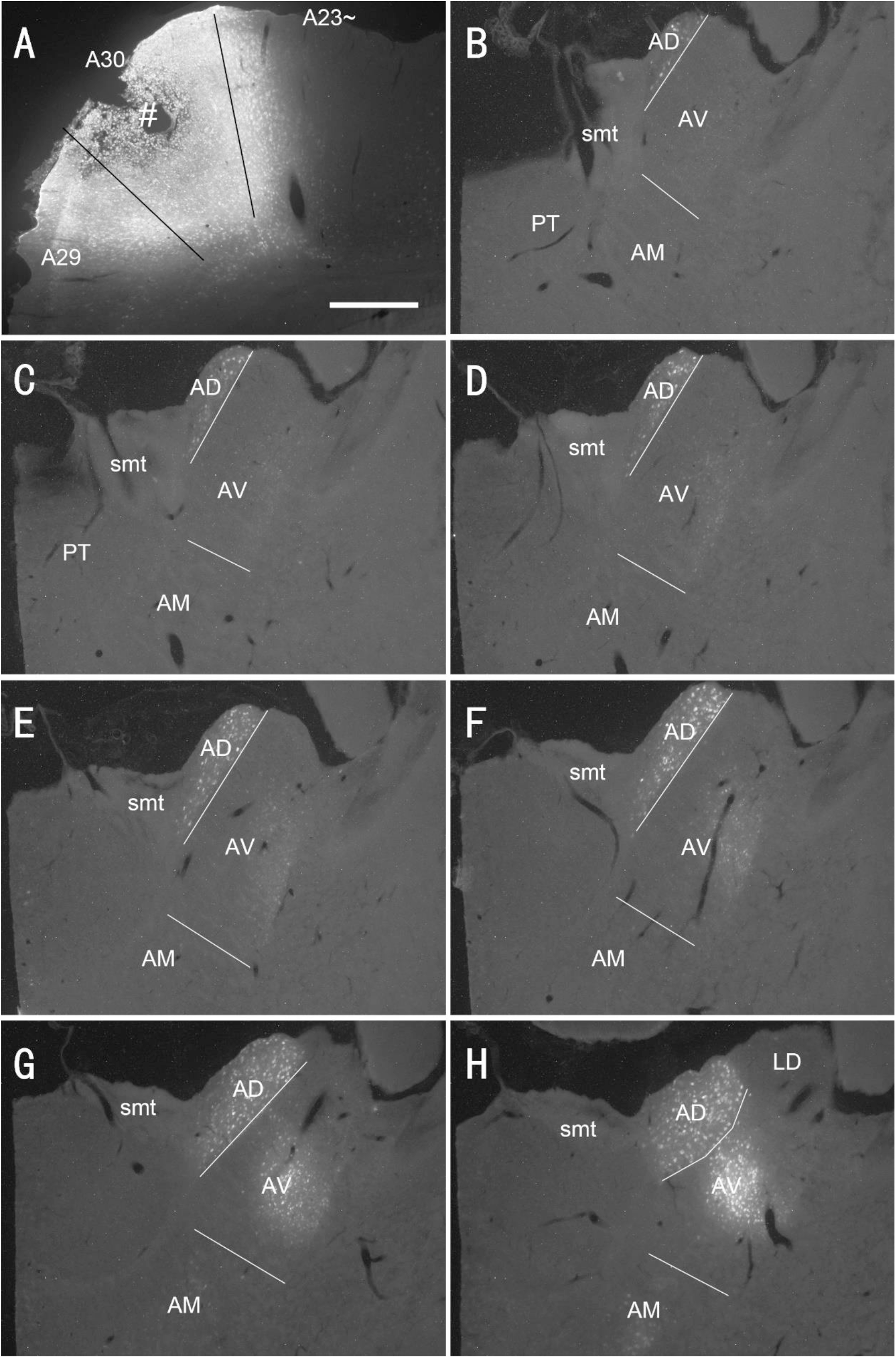
Labeled neurons in the AD and AV following FG injection in the rat A30. (A). One FG injection site (#) in A30 with slight involvement in A29. (B-H) Sequential coronal sections from the anterior (B) to posterior (H) levels showing FG labeled neurons in the AD, AV and AM. The borders between the AD and AV and between AV and AM are indicated by a solid line and a dashed line, respectively. Note that most of the labeled neurons are observed in both AD and AV with more at the caudal levels (E-H). A few of labeled neurons are also detected in the posterior AM (H). Bar: 500 μm in (A) for all panels.

Quantitative analysis of the labeled neurons in the AD, AV and AM following FG injections in A23~ or A30 further indicates that A23~ but not A30 receives major inputs from the AM (Fig. 4A, left) (F [2, 27]=17.42, p<0.0001, [R-squared: 0.7597]) whereas A30 but not A23~ receives inputs from both the AD and AV (Fig. 4A, right) (F [2, 27]=7.833, P<0.001, [R-squared: 0.4885]). Comparison of the labeled neurons resulted from FG injections in A23~ and A30 has further confirmed the differences (p<0.0001; see Fig. 4B). It is also noted that FG Injections in posterior A23~ lead to labeled neurons across the A-P extent of the AM with more in the middle levels (e.g., Fig. 4C) while the injections in posterior A30 produce labeled neurons mainly in the posterior AD and AV (e.g., Fig. 4D).

**Figure 4.**
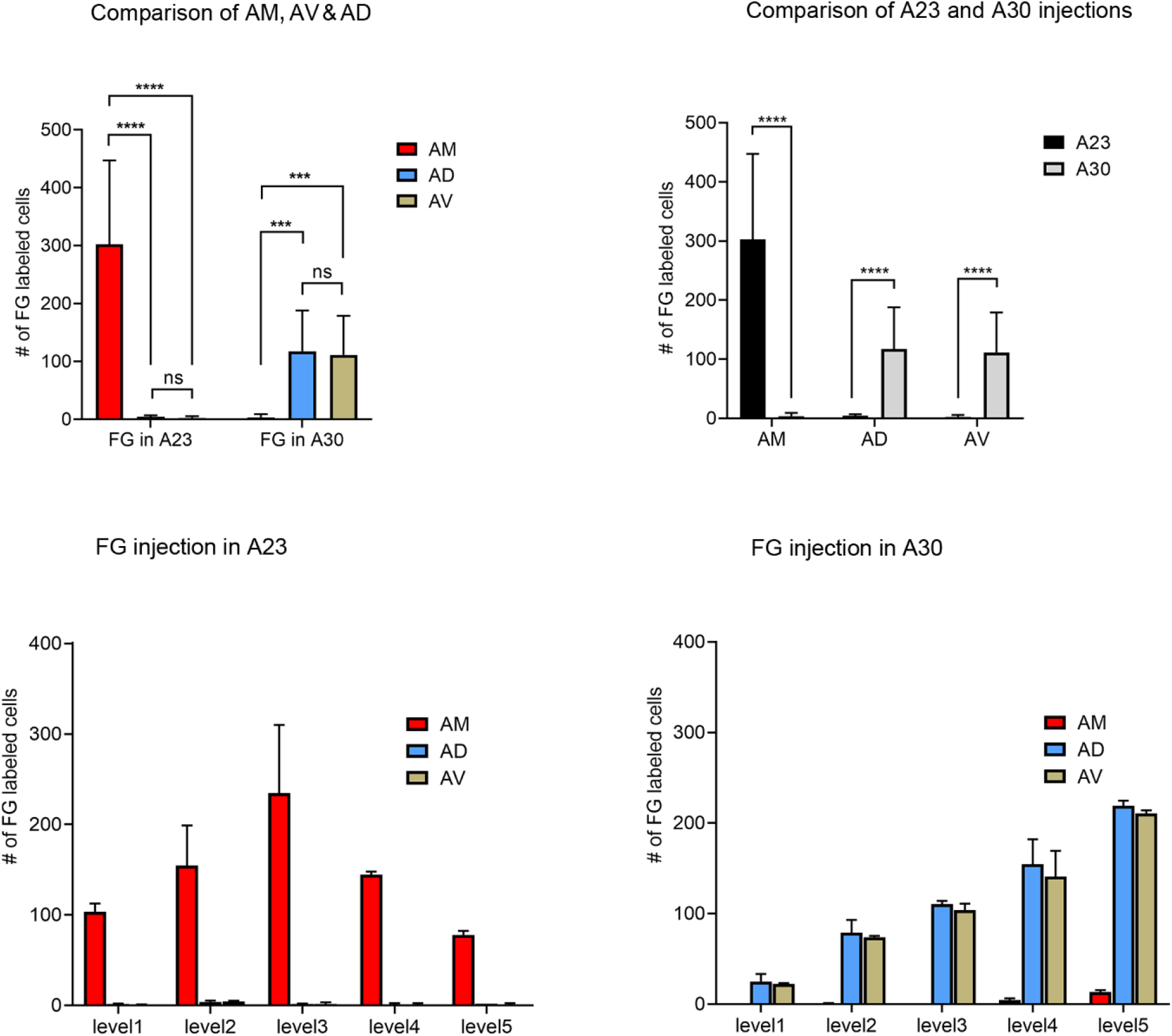
Quantitative analysis of FG labeled neurons in the AD, AV, and AM of the rats. (A) Comparison of the cell counts in the AM, AD and AV following A23~ (left) and A30 (right) FG injections. ***P<0.001; ****P<0.0001. (B) Comparison of the cell counts in each of the anterior thalamic nucleus between A23~ and A30 injections. ****P<0.0001. (C) Example of the cell counts along the A-P extent in one case with an FG injection in posterior A23~ (mean ± SD). (D) Example of the cell counts along the A-P extent in one case with an FG injection in posterior A30 (mean ± SD).

Based on above findings it is reasonable to infer that anterograde tracer injections in the AM would produce obvious terminal labeling in A23~ rather than in adjoining A30, and thus enable identification of the border between A23~ and A30. Indeed, following the BDA injections into the AM of the rats (4 cases), BDA labeled axon terminals were observed across the A-P extent of A23~. For example, one small BDA injection in the dorsomedial part of the middle AM resulted in terminal labeling across the A-P extent of A23~ with relatively densely and weakly labeled terminals in the anterior and posterior portion of A23~, respectively (data not shown). In contrast, following one larger BDA injection covering most of the AM (Fig. 5A), the resulted terminal labeling is clearly observed in almost all A-P extent of A23~ (Fig. 5B-I). Labeled terminals are distributed mainly in layers 1, 5 and 6 of A23~ (Fig. 5J). The labeled axon terminals in layer 1 of A23~ extend slightly into A30 (medially) and the medial visual cortex (V2M; laterally) (see the middle column of Fig. 5) while those in layers 5 and 6 mostly concentrate in A23~. This labeling pattern enables the identification of the medial and lateral borders of A23~. The anterior and posterior borders of A23~ can also be determined based on the A-P extent of the densely labeled terminals resulted from the larger BDA injection (Fig. 5A-I). It is estimated that A23~ in the rats corresponds roughly to the RSagl defined by Swanson (Swanson, 2004) or medial portion of the V2M (V2MM) defined by Paxinos and Watson (2013). Specifically, rat A23~ anteriorly starts at the level above the CCS (the level slightly rostral to Fig. 5B), and c posteriorly ends at the level, where the terminals are located at the medial edge of the visual cortex (at the level of Fig. 5I; about 8.70mm caudal to the bregma based on the Paxinos atlas).

**Figure 5.**
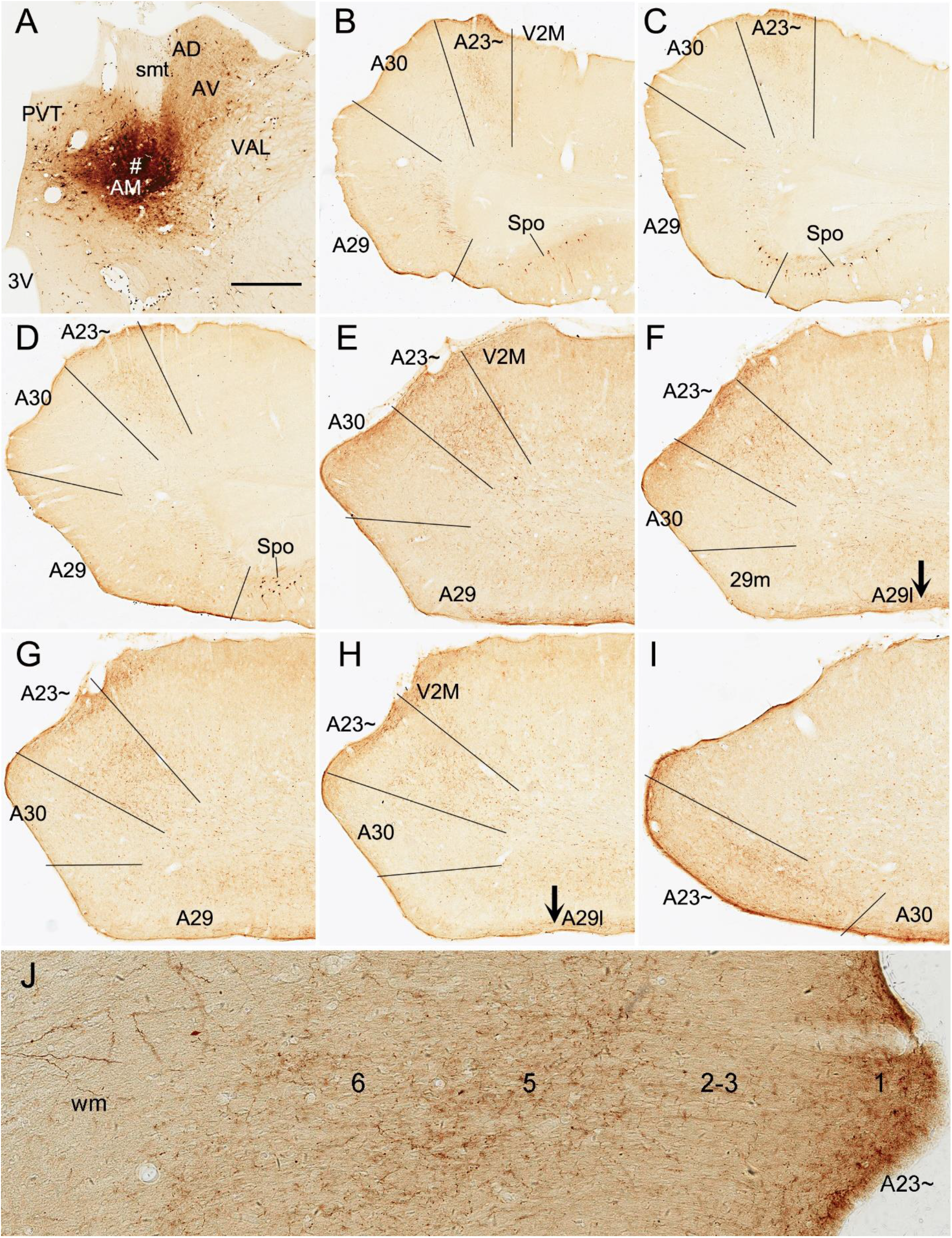
AM projections to A23~ in the rat, revealed with BDA. (A) One BDA injection site (#) in the AM. (B-I) Sequential coronal sections from the anterior (B) to posterior (I) levels showing BDA labeled axon terminals in A23~. The regional borders are indicated by solid lines in each panel. Labeled terminals are mainly detected in layers 1, 5 and 6 of A23~ and in layer 1 of A29l (indicated by the arrows in F and H). Labeled terminals in layer 1 of A23~ also extend into layer 1 of A30 and V2M for a short distance. Note that some BDA labeled neurons are observed in the polymorphic layer of the subiculum (Spo in B-D). Note also that A23~ shifts slightly from lateral to medial positions along the A-P axis. (J) Higher magnification view of the labeled axon terminals in A23~ shown in (B). Bar: 500 μm in (A) for all panels.

### 3.3 Anteromedial thalamic inputs define A23~ in mice

To confirm above findings in mice, we performed a survey on the Allen Mouse connectivity dataset (https://connectivity.brain-map.org), in which more sensitive anterograde viral tracers were used for the experiment of both wild-type (2 cases) and Cre-line (3 cases) mice. As in the rats, the AM injections produced dense terminal labeling in A23~ rather than in adjoining A30. For instance, one injection of the viral tracers into the AM of the mouse (Figs. 6A; 7A) results in dense terminal labeling in A23~ with no or few in adjoining A30 (Fig. 6B-L). Like in the rats, labeled axon terminals in A23~ mainly concentrate in layers 1, 5 and 6 (Fig. 6B-L). It is noted that layer 1 of the lateral A29 (A29l or A29a) also contains dense terminal labeling but this labeling is far away from that in A23~ (separated by A30 and medial A29: A29m or A29bc). In contrast to the AM injections, viral tracer injections in the AV (e.g., Fig. 6M) lead to heavy terminal labeling in layer 1 and moderate labeling in layers 3, 5-6 of A29m (Fig. 6N-P) with much less in A30 (Fig. 6N) and no in A23~ (Fig. 6N-P). It is noted that the AV but not AM injections also produce axon terminal labeling in the polymorphic layer of the subiculum (Spo; Fig. 6P; also see Ding et al., 2020). The overall brain-wide projection pattern resulted from the AM injection in Figure 6A are shown in the lateral and dorsal aspects of the brain (see Fig. 7A and 7B, respectively). It is obvious that five major target regions of the AM projections are the anterior cingulate cortex [ACC, including the anterior cingulate area (ACA) and prelimbic area (PL)], CPu (caudate-putamen), PrS, A29l and A23~ (Fig. 7A, B). The moderately labeled terminals in the infralimbic cortex (IL), nucleus accumbens (NAC) and basolateral nucleus of amygdala (BL) are probably originated from the involvement of the injection in the parataenial nucleus (PT), which was reported to project to these targets (Vertes and Hoover, 2008). It is also clear from the dorsal view of the brain that A23~ is a band-like terminal zone (outlined in Fig. 7B) located lateral to A30, which does not receive significant inputs from the AM (see Fig. 6B-I).

**Figure 6.**
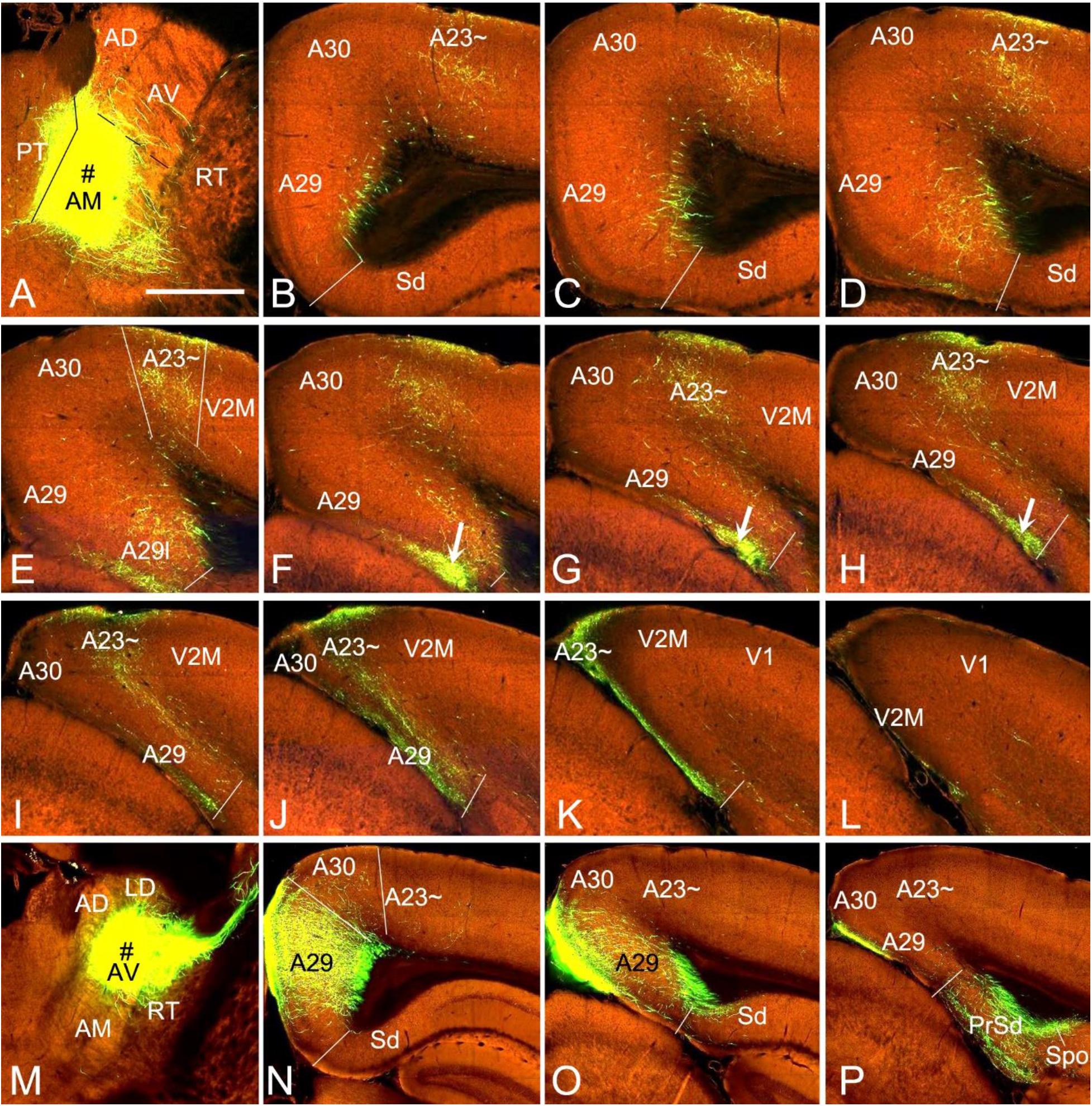
AM and AV projections to A23~ and A29 in the mouse, revealed with viral tracers. (A) One viral injection site (#) in the AM with slight involvement in the paratenial nucleus (PT). (B-L) Sequential coronal sections from the anterior (B) to posterior (L) levels showing labeled axon terminals in A23~ and area 29l (arrows in F-H) resulted from the AM injection. Like in the rat, labeled terminals are mainly detected in layers 1, 5 and 6 of A23~ and layer 1 of A29l. (M-P) One viral tracer injection site (# in M) centered in the AV of the mouse results in labeled axon terminals in layers 1, 5 and 6 of A29 with much fewer in A30 and none in A23~ (N, O) and in the PrSd and polymorphic layer of the subiculum (Spo; P). Bar: 560 μm in (A) for all panels.

**Figure 7.**
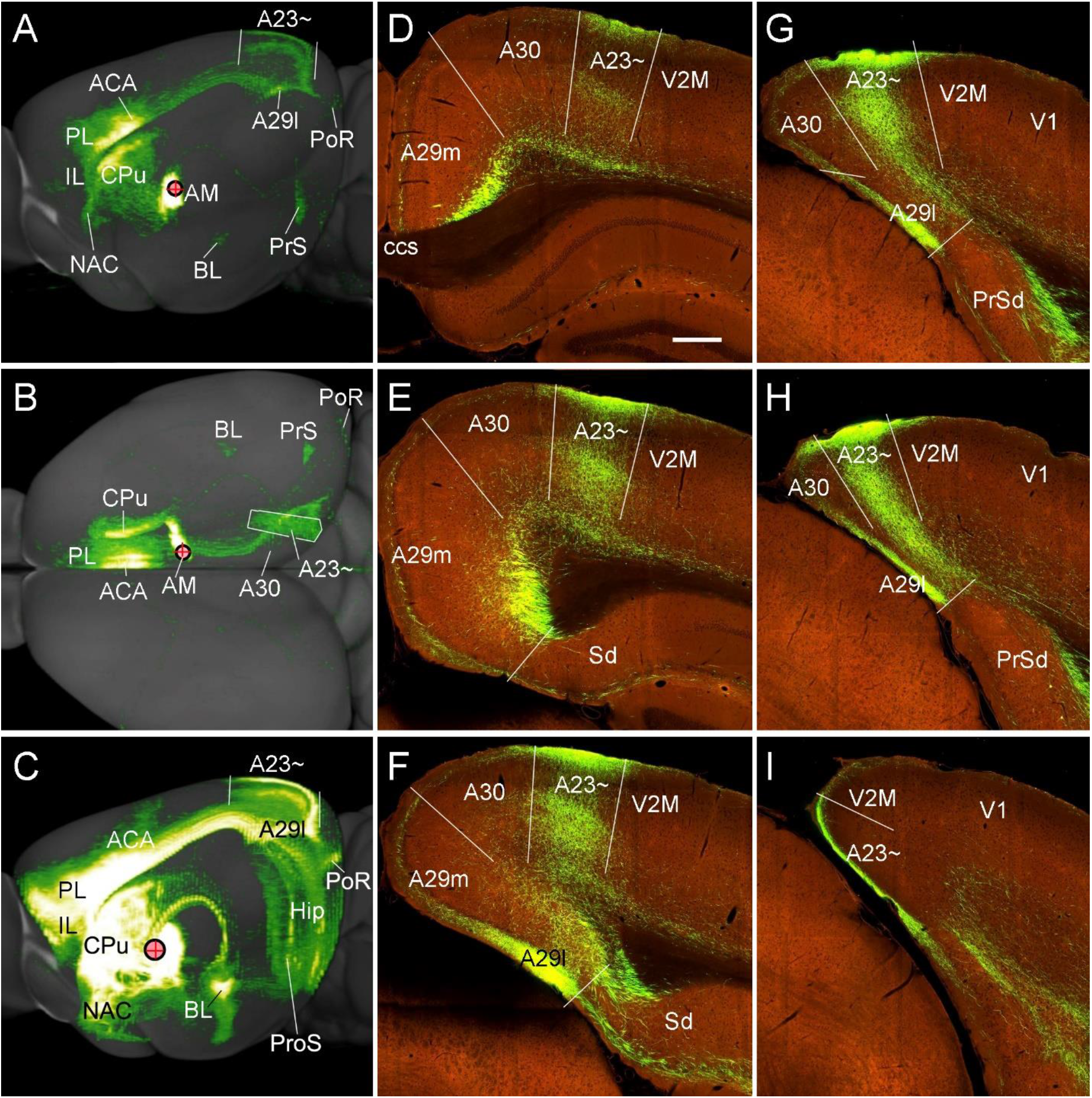
Major targets of the AM projections in the mice, revealed with viral tracers. (A, B) The AM projection pattern on the lateral (A) and dorsal (B) aspects of the same brain. Individual A-P sections showing the labeled terminals are demonstrated in Figure 6A-L. It is obvious that the major targets of the AM projections include the ACA, PL, A23~, A29l, CPu and PrS. Note that the injection is slightly involved in the PT, which leads to some terminal labeling in the IL, NAC and BL. (C) One viral tracer injection site covering entire AM and adjoining PT and reuniens nucleus (Re) results in strong terminal labeling in A23~ but not in A30. (D-I) Sequential coronal sections showing the A-P extent of the mouse A23~ revealed by the injection shown in (C). The strongly labeled axon terminals in A23~ make this region easily identifiable. Generally, A23~ starts anteriorly at the level overlying the CCS (D) and ends posteriorly at the medial edge of the posterior cortex (slightly posterior to the level H. Bar: 280 μm in (D) for panels (D-I).

To determine and verify the full A-P extent of A23~ in the mice, we also examined some cases with even larger injections which cover full A-P extent of the AM as well as the adjoining PT and reuniens nucleus (Re) of the thalamus (3 cases). For example, as shown in Figure 7C, although strong terminal labeling is found in many other structures the A-P extent of the labeled terminals in A23~ is comparable to that revealed in Figure 7A. Specifically, the most anterior level of A23~ (containing densely labeled terminals from the AM) is at the level above the CCS (Fig. 7D) while the most posterior level is at the level between panels (H) and (I) of Figure 7 (about 4.36mm caudal to the bregma according to the mouse brain atlas (Paxinos and Franklin, 2012). It is noted that the location of A23~ slightly shifts from lateral (Fig. 7D) to medial (Fig. 7H) positions along the A-P axis (Fig. 7D-I). It should also be mentioned that the PT and Re do not project to A30 and A23~ (Vertes and Hoover, 2008) although these two regions are involved in the large injection site. The PT mainly projects to the IL, NAC, and BL (Vertes and Hoover, 2008) while the Re mostly projects to the prosubiculum (ProS in Fig. 7C; also see Figure S9 of Ding et al., 2020), CA1, piriform, insular, perirhinal (PRC) and prefrontal cortices (Vertes et al., 2006).

### 3.4 Afferent and efferent connections of A23~ in the rats

To explore whether connections of A23~ in the rodents are comparable to those of A23 in the monkeys, we investigated brain-wide distribution of the retrogradely labeled neurons following FG injections in A23~ of the rats (4 cases). For instance, as shown in Figure 8, after one FG injection into A23~ (Fig. 8A), a variety number of the labeled neurons are seen in cortical A29 (layers 3, 5; Fig. 8B), medial orbital cortex (ORBm, layers 2-3; Fig. 8C), ACA (posterior area 24, layers 3, 5; Fig. 8D) and parietal association cortex (PtA). Sparsely labeled neurons are also detected in layer 5 of the secondary motor cortex (M2; Fig. 8D), PL, postrhinal cortex (PoR), medial entorhinal cortex (MEC), a part of lateral secondary visual cortex (V2L) and the posteior part of the temporal association cortex (TeA) located posterior to the primary auditory cortex. The subcortical regions containing the labeled neurons include the AM (Fig. 8E), laterodorsal thalamic nucleus (LD; Fig. 8F), claustrum (Cla; Fig. 8G),_nucleus coeruleus (NC; Fig. 8H), forebrain basal nucleus and full A-P extent of the medial part of the lateroposterior nucleus-pulvinar complex (LP-Pul; Fig. 9A, C, E and G), which appears to correspond to the medial pulvinar (Pulm) of the monkeys (Baleydier and Mauguiere, 1985).

**Figure 8.**
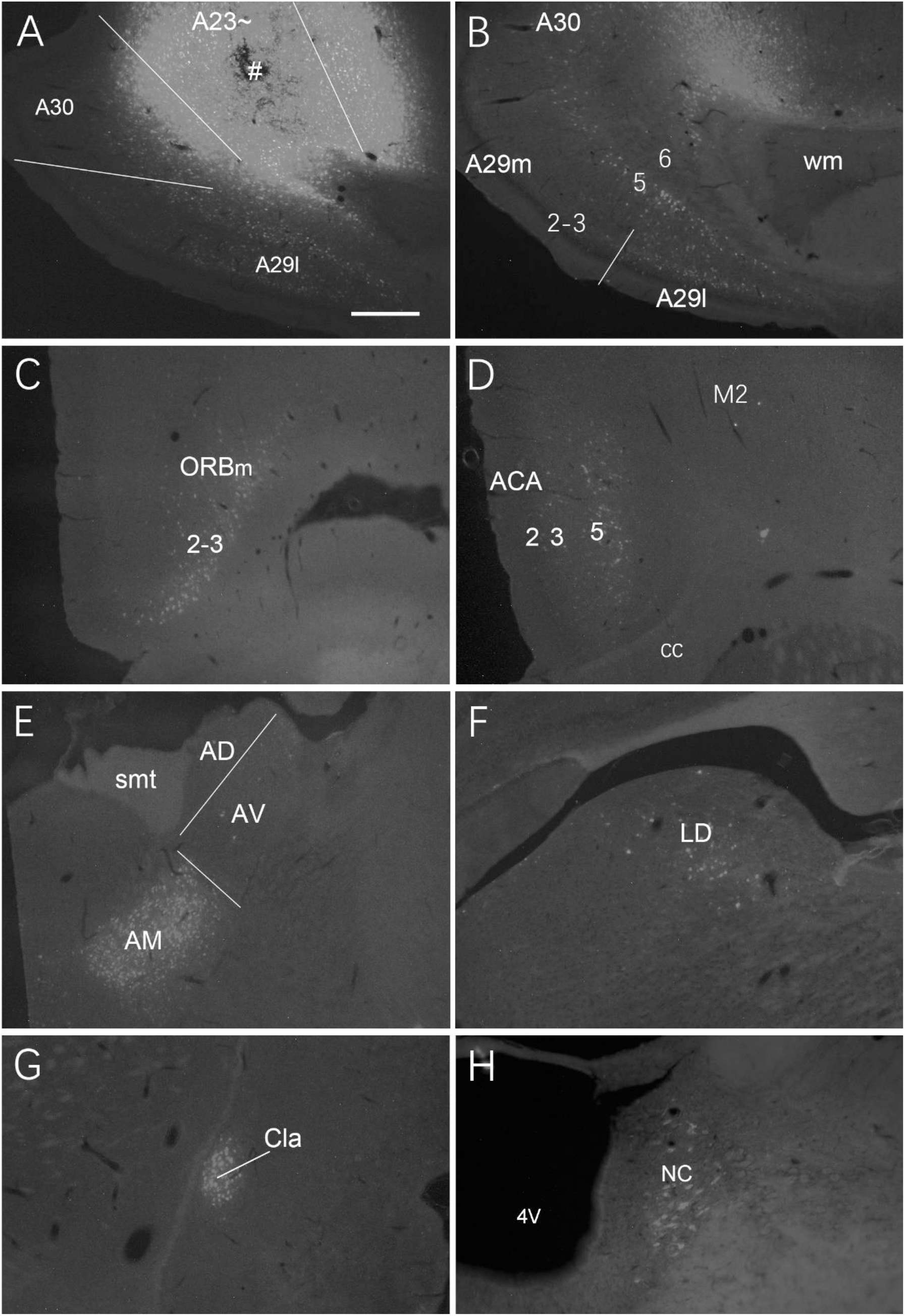
Afferent projections of A23~ in the rat, revealed with FG. (A) One FG injection site (#) in A23~. (B-H) Examples of FG labeled neurons in A29 (B), ORBm (C), ACA and M2 (D), AM (E), LD (F), Cla (G) and nucleus coeruleus (NC; H). Bar: 500 μm in (A) for all panels.

**Figure 9.**
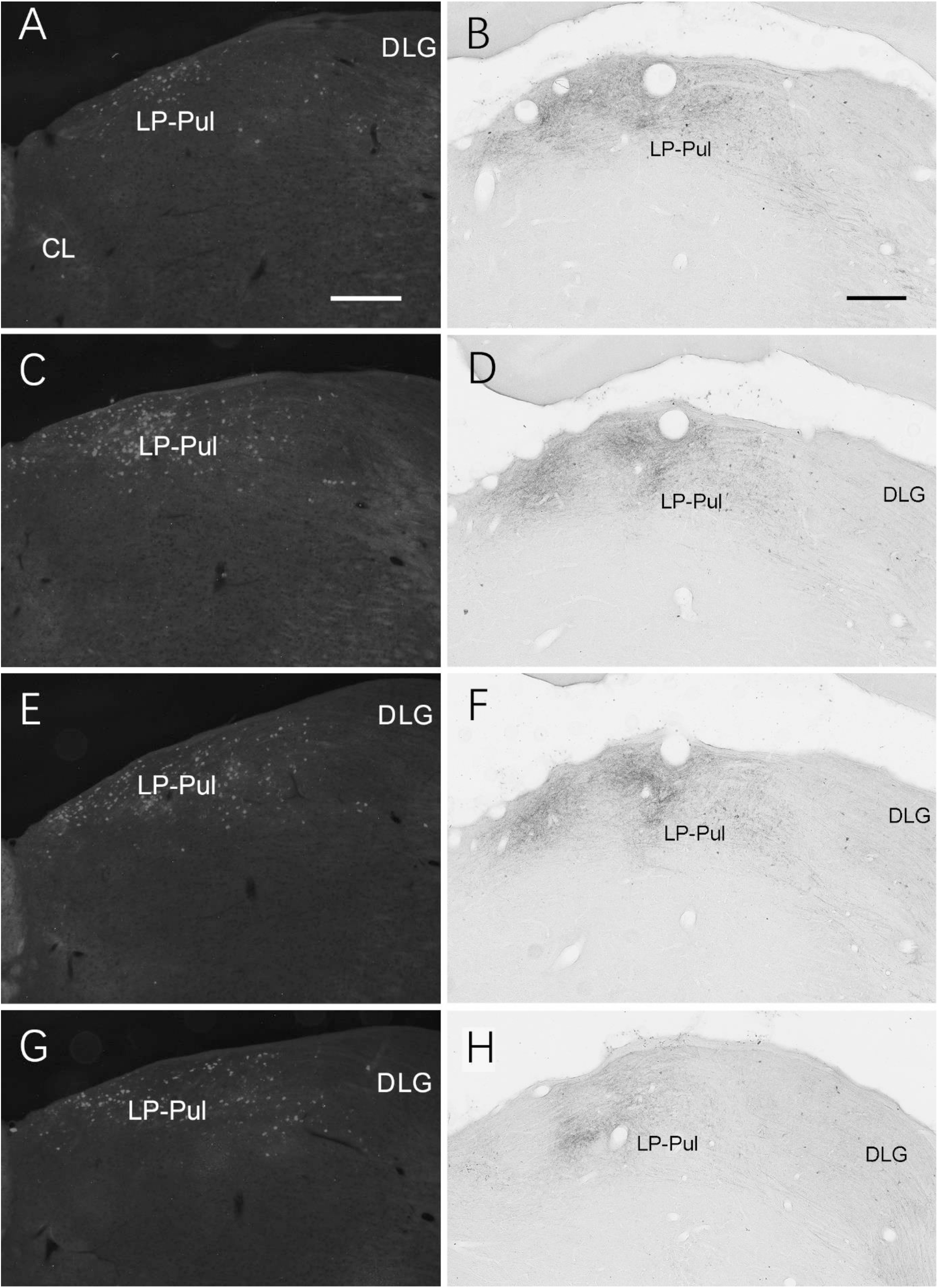
Labeled neurons and axon terminals in the LP-Pul of the rats. (A, C, E and G) Sequential coronal sections from the anterior (A) to posterior (G) levels of the LP-Pul showing the labeled neurons in the medial part of the LP-Pul after FG injection in A23~. (B, D, F and H) Sequential coronal sections from the anterior (B) to posterior (H) levels of the LP-Pul showing the labeled axon terminals in the LP-Pul after the BDA injection in A23~. Note that more terminals are seen in the medial part of the LP-Pul. Bars: 500 μm in (A) for panels (A, C, E and G); 300 μm in (B) for panels (B, D, F and H).

Brain-wide distribution of anterogradely labeled axon terminals was also examined following BDA injections into A23~ of the rats (4 cases). For example, one BDA injection restricted in A23~ (Fig. 10A) leads to terminal labeling mainly in layers 1 and 5 of ORBm (Fig.10B), layers 1 and 3 of ACA (Fig.10C, C’), AM (Fig.10D), Cla (Fig. 10E), LD (Fig. 10F), layer 3 of the PrS-PoS (Fig.10G), and layers 5-6 of the PoR (Fig. 10H). Interestingly, like the distribution of the neurons projecting to A23~, full A-P extent of the medial part of the LP-Pul receives strong inputs from A23~ (e.g., Fig. 9B, D, F and H). Weak terminal labeling is also detected in layers 5-6 of the V2L (Fig. 10I), layer 1 of the PL and in the M2, MEC, pretectal nucleus (PTN), superior colliculus (SC), dorsomedial CPu, zona incerta (ZI), anterolateral portion of the pontine nucleus (PN) and the PtA.

**Figure 10.**
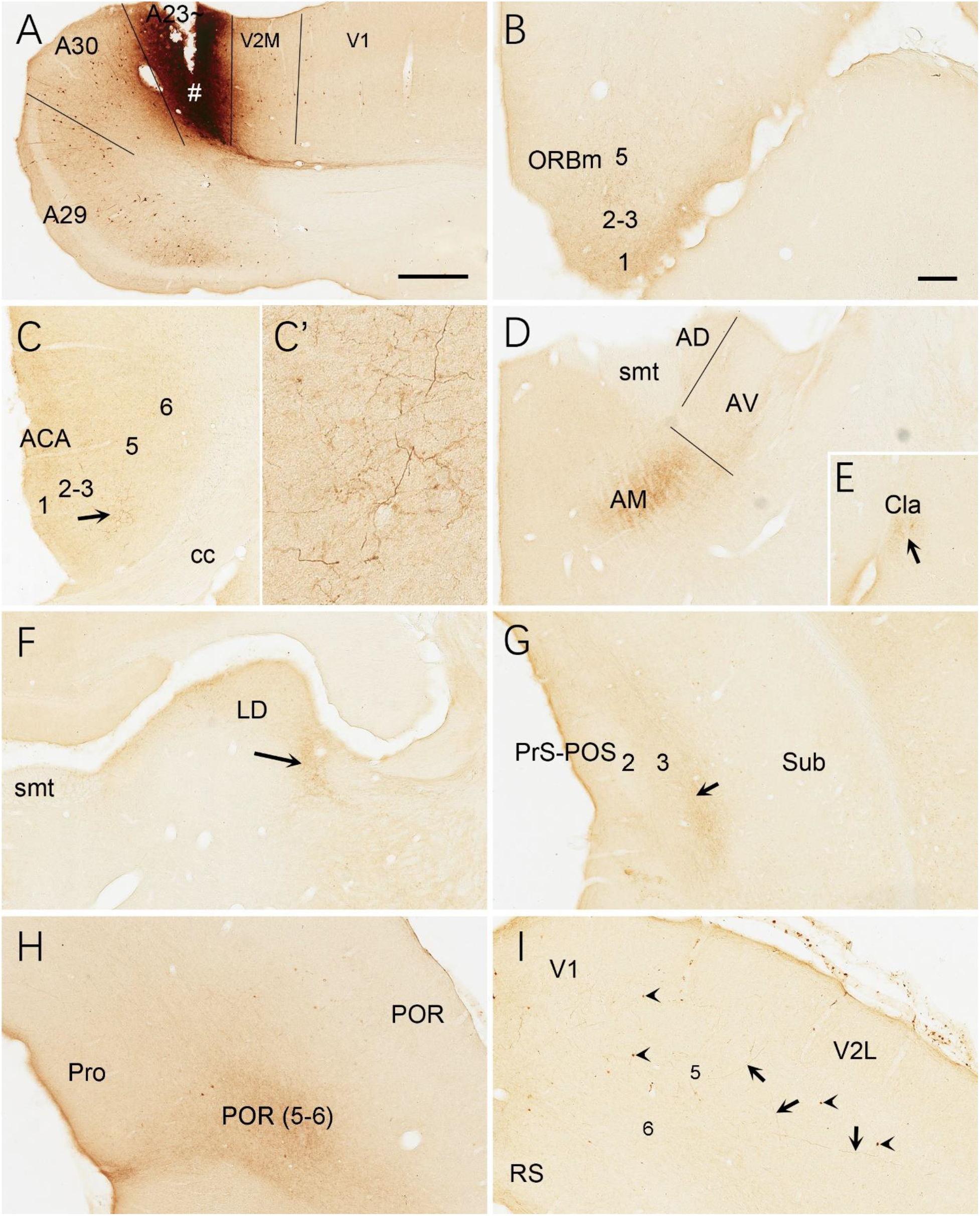
Efferent projections of A23~ in the rat, revealed with BDA. (A) One BDA injection site (#) restricted in A23~. (B-I) BDA labeled axon terminals in layer 1 of the ORBm (B), layers 1 and 3 of the ACA (C and C’), AM (D), Cla (E), LD (F), layer 3 of the PrS-PoS, layers 5-6 of the PoR (H) and V2L (arrows in I). The arrows in (C, E, F and G) point to the regions with dense axon terminals. A high magnification view of the terminals in the ACA (arrowed region) is shown in (C’). Note that many BDA labeled neurons are observed in layer 5 of A29, A30 and V2M (A) with a few in V1 (A) and V2L (arrowheads in G). A few of BDA labeled axon terminals are also found in layer 1 of the PL-IL, layer 6 of the MEC and the dorsomedial part of the CPu (not shown). Bars: 500 μm in (A) for panels (A, C and G); 300 μm in (B) for panels (B, E, H and I).

### 3.5 Efferent projections of A23~ in the mice

To further confirm the efferent projections of A23~ using a more sensitive tracing method we searched the Allen connectivity dataset for the cases with viral tracer injections in the A23~ (5 cases). For example, as shown in Figure 11, the injection in the anterior A23~ of the mouse (see injection site in Fig. 11A-C, G) produces strong axon terminal labeling in both ipsi- and contra-lateral A23~, A29l and AM (Fig. 11A, C) and many ipsilateral structures (Fig. 11A-K). The ipsilateral structures include the ORBm (Fig. 11B, C, D), PL and CPu (Fig. 11B, C, E), V2L (Fig. 11A, C, F), A29, VLG-m and ZI (Fig. 11A-C, G), LP-Pul (Fig. 11B, G), ACA and M2 (Fig. 11B, C, H), LD and PtA (Fig. 11B, I), V2M, posterior TeA and A23~ (Fig.11C, J), PoR and MEC (Fig. 11C, K), nucleus of lateral lemniscus (NLL), SC and PAG (Fig. 11B, C, K). Labeled axon terminals are also seen in the posterior CPu (CPu-p), anterolateral pontine nucleus (PN), Cla, PrS-PoS, and PaS (Fig. 11A-C) as well as in the deep layers of the PRC, posterior hypothalamic nucleus (PHN), ventral anterior thalamic nucleus (VA), VM, lateral dorsal tegmental area (LDTg), pontine reticular formation (PnRF), paramedial raphe nucleus (PMnR) and midline thalamic nuclei (MiN) such as the central lateral nucleus of the thalamus (CL) and Re.

**Figure 11.**
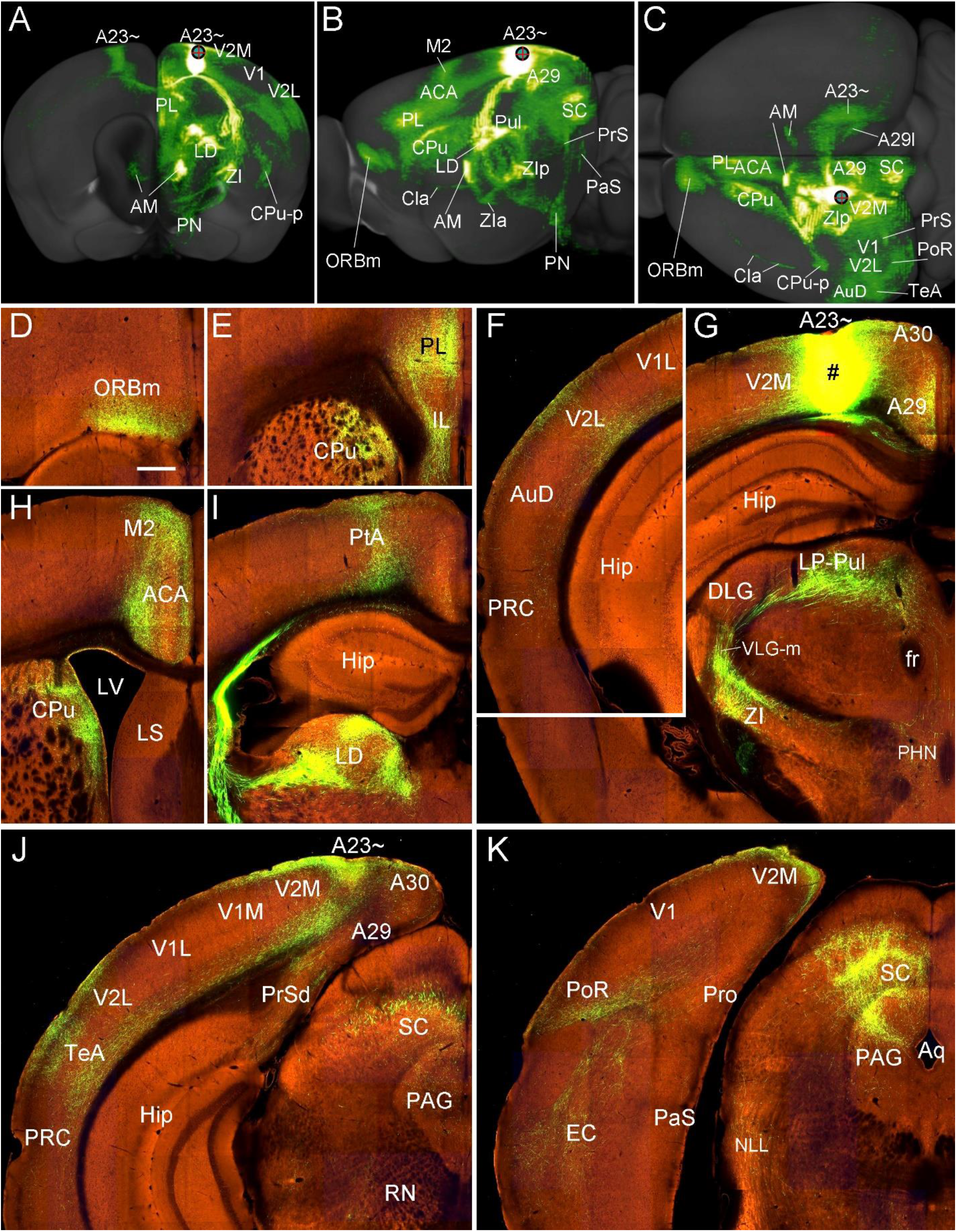
Efferent projections of A23~ in the mouse, revealed with viral tracers. (A-C) Overall projection pattern of the A23 projections in a *Emx1*-Cre mouse on the anterior (A), lateral (B) and dorsal (C) aspects of the brain. Some major target regions are marked. Note the obvious terminal labeling in the ipsi- and contralateral AM and A23~ (A and C). (D-I). Axon terminal labeling in the ORBm (D), PL, CPu (E, H), V2L (F), LP-Pul, ZI (G), ACA, M2 (H), PtA, LD (I), posterior TeA and A23~ (J), PoR, EC, SC and PAG (K). For orientation of the sections in panels (D-K), medial is at the right and dorsal is at the top. Bar: 400 μm in (D) for panels (D-K).

### 3.6 Brain-wide connections of A30 in the rats

As a comparison with A23~, brain-wide connections of A30 were also examined in the rats (3 cases). For instance, following one BDA injection in A30, some retrogradely labeled neurons are found to distribute in layer 5 of A29, V2M, V1 and ACA (Fig. 12A, B), AD and AV (Fig. 12C), LD (Fig. 12D), LP-Pul (Fig. 12E), posterior A30 and posterior V2M (Fig. 12G), Cla, V2L and M2. As seen in the same case, BDA labeled axon terminals are detected in A29 (layers 1, 5; Fig. 12A), ACA (layers 1, 3; Fig. 12B), M2 (layers 1, 2-3; Fig. 12B), AV (Fig. 12C), LD and RT (Fig. 12D), LP-Pul (Fig. 12E), PrS-PoS (layers 1, 3; Fig. 12F), posterior V2M and A30 (layers 2-3; Fig. 12G), layer 1 of the PaS and layers 5-6 of the PoR (weak labeling in Fig. 12H), layers 1 and 6 of the V1 and V2L (weak labeling), SC, PTN, ZI, PN, and CPu. Compared to BDA injections, FG injections in A30 labeled much more neurons in the regions containing BDA labeled neurons.

**Figure 12.**
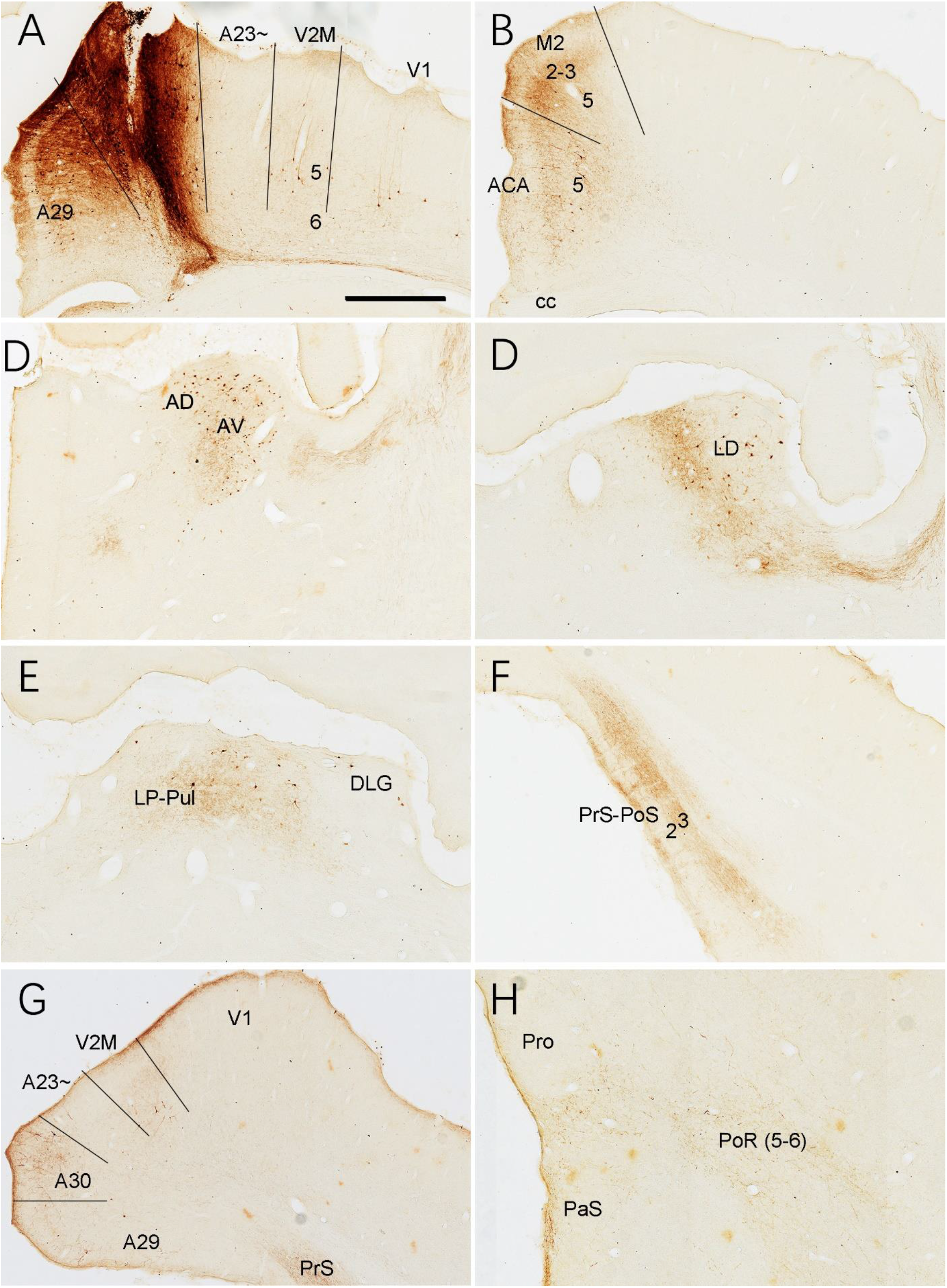
Connections of A30 in the rat, revealed with BDA. (A) One BDA injection site (#) in A30. Note that BDA labeled neurons are observed in A29, V2M, V1 (A). (B-H) BDA labeled neurons and axon terminals in the representative regions. Note the BDA labeled neurons in ACA (B), AD, AV (C), LD (D), LP-Pul (E), V2M and the caudal A30 (G). BDA labeled axon terminals are detected in A29 (A), ACA, M2 (B), AV (C), LD (D), LP-Pul (E), PrS-PoS (F), V2M, posterior A30 (G), PaS and PoR (H). Bar: 300 μm in (A) for all panels.

## 4. Discussion

Since Brodmann’s comparative cortical mapping, the PCC (mainly A23) has been treated as a unique structure in human and NHP brains. Lower animals such as rabbits and rodents were not reported to have the equivalent of the primate PCC based on cytoarchitectonic criteria (Brodmann, 1909; Swanson, 2004; Vogt et al., 2005; Sugar et al., 2011; Wang et al., 2020). However, functional studies suggest that the PCC is an important component of the default mode network (DMN), which is reported in both primates and rodents (Mantini et al., 2011; Lu et al., 2012; Stafform et al., 2014; Hsu et al., 2016). The DMN (including the PCC) in human brains is heavily involved in autobiographical information, self-reference, thinking about others, spatial episodic memory, and plan for future (Andrews-Hanna, 2012; Leech and Sharp, 2014; Rolls, 2019). These functions are critical to survival and thus the PCC (mainly A23) should also exist in the rodents. The present study has revealed the rodent equivalent of the primate A23 in both rats and mice based on topographical relationship with A30 and conservative AM inputs to A23 as well as the differential expression patterns of the genes *Tshz2, Penk, Cux2* and *Igfbp5* in the RSagl (corresponding to A23~ in this study) from A30 of the mice (Hu et al. 2020; Lu et al., 2020). Since we can not distinguish A23 from A31, and A31 is probably very small in the rodents (if it exists), we have termed the rodent equivalent of the PCC as A23~ in this study. The present study has also shown that the afferent and efferent connections of A23~ in the rodents are generally comparable to those of the primate A23 and that the rodent A23~ has differential connections from the retrosplenial A29 and A30. Finally, our findings also suggest that the feature of strong AM innervation of A23 could be used to consistently distinguish A23 from adjoining A30, V2M and medial PtA across species.

## 4.1 Anteromedial thalamic inputs define rodent cingulate cortical area 23

The present study has defined the location and extent of the rodent equivalent of the primate A23 (i.e., A23~). Like in the primates, A23~ in both rats and mice receives its main inputs from the AM rather than from the AD and AV. In contrast, the adjoining A30 (RSag) receives its main inputs from the AD and AV rather than from the AM (see Fig. 4). Therefore, following an anterograde tracer injection covering all or almost all extent of the AM, the labeled axon terminal field located lateral to A30 (RSag) would mark the location and extent of A23~. In this study, the terminal field in A23~ identified with the AM inputs roughly occupies the region previously treated as retrosplenial area 29d (Shibata, 1993), RSagl (Swanson, 2004), a part of the RSag (Sripanidkulchai and Wyss, 1986; Wyss and van Groen, 1992) or as the visual area V2MM (Paxinos and Franklin, 2012; Paxinos and Watson, 2013) or area 18b (Miller and Vogt, 1984; van Groen et al., 1999). Specifically, in coronal sections, A23~ in the rats and mice forms an A-P band starting anteriorly at the level overlying the CCS and ending posteriorly at about the level where the AM inputs are located at the medial edge of the posterior cortex (e.g., Fig. 7D-I). This finding of AM projections to A23 (but not to A30) is consistent with those reported in primates (Baleydier and Mauguiere; 1980; Vogt et al., 1987; Shibata and Yukie, 2003). For example, when the retrograde tracers were restricted to A23 of the monkeys, the labeled neurons were mostly observed in the AM with no or few in the AD and AV (Baleydier and Mauguiere; 1980; Vogt et al., 1987; see also cases M-2-WGA-HRP and M-5-CTb in Shibata and Yukie, 2003). However, when the tracers were involved in adjoining A30, labeled neurons could also be found in the AD and AV in addition to the AM (see case M1-WGA-HRP in Shibata and Yukie, 2003; cases 1 and 2 in Buckwalter et al., 2008). On the other hand, when the tracers were restricted in A30 and A29 of the monkeys, many labeled neurons were seen in the AD and AV with no or few in A23 (see cases M3-CTb and M5-DY in Shibata and Yukie, 2003).

## 4.2 Comparative connectivity of area 23 in primates and rodents

The relative size of A23 in NHP and human brains is much larger compared to A29 and this A23 can be further divided into three subdivisions (A23a, b and c) (see Fig. 1A, B for monkey and Ding et al., 2016 for human). In sharp contrast, the relative size of A23~ in the rodents is much smaller in comparison with A29. However, the major afferent and efferent connections of A23 in the rodents and primates appear similar and comparable. For example, A23 in the primates is reported to receive afferents from the AM, Pulm, Cla, dorsolateral frontal cortex (DFC; including areas 9, 10 and 46), ORBm (areas 11,13 and 14), ACC (A24 and A32), retrosplenial cortex (A29 and A30), posterior parahippocampal cortex (PHC; including areas TH, TL and TF), posteromedial inferior parietal cortex (areas Opt and PGm), posterior superior temporal cortex (area TAc or Tpt) and dorsal bank of the superior temporal sulcus (area TPO) (Baleydier and Mauquiere, 1980, 1985; Vogt and Pandaya, 1987; Vogt et al., 1987; van Hoesen et al, 1993; Yukie, 1995; Kobayashi and Amaral, 2003; Shibata and Yukie, 2003; Buckwalter et al, 2008; Seltzer and Pandya, 2009). The efferent targets of A23 in the monkeys mainly include the AM, SC, PTN, PN, Cla, LD, LP, Pulm, CPu, PrS, A29-30, ACC, PHC, ORBm, DFC, areas TPO and TAc/Tpt (Pandya et al, 1981; van Hoesen et al., 1993; Shibata and Yukie, 2003; Parvizi et al, 2006; Kobayashi and Amaral, 2007; Seltzer and Pandya, 2009). Similarly, the rat A23~ receives its main inputs from the AM, medial LP-Pul (similar to the monkey Pulm), Cla, a part of M2 (likely similar to A24c in the monkeys), ORBm, ACA (similar to A24a and b in the monkeys), A29-30, PoR (similar to the monkey PHC), medial PtA (similar to area Opt/PGm in the monkeys), a part of the V2L (similar to area TPO in the monkeys) and the posterior TeA (similar to the monkey area TAc/Tpt). The A23~ in the rats and mice mainly projects to the AM, SC, PTN, PN, Cla, LD, LP-Pul, CPu, PrS, A29-30, ACA, PoR, ORBm, a part of M2, a part of the V2L and the posterior part of the TeA (e.g., Fig. 11). However, the rodent equivalent of the monkey DFC (areas 8, 9, 10, 46) remains to be identified.

## 4.3 Comparison of area 23~ with areas 30 and 29 in rodents

As mentioned above, A29 in the rodents is easily distinguishable from A30 due to its unique outer granular layers 2-3. However, the border between rodent A30 and A23~ is difficult to be identified, particularly in Nissl-stained sections since rodent A23~ does not display a thin inner granular layer 4 while the primate A23 does. This is the major reason why A23~ was not identified in previous studies. However, given the primate cortex having overall much more granular cells in layer 4 compared to the rodent cortex, it is not so surprising to see few granular cells in layer 4 of the rodent A23~. In literature, significant differences in cytoarchitecture are commonly observed between the primate and rodent brains. For example, hippocampal CA1 cells in NHP and human brains are lightly stained and loosely packed while those in the rat and mouse brains are darkly stained and very densely packed (see Ding, 2013). However, the topographic relationship of CA1 with CA2/CA3 and ProS/Sub as well as the connectional patterns of these hippocampal subfields remain similar across species (e.g., Ding et al., 2020). The present study has provided a connectional approach that enables the differentiation of A30 and A23. The AM inputs predominantly terminate in A23~ with few in adjoining A30 and V2M while the inputs from the AD and AV mostly innervate A29 with fewer in A30 and almost no in A23~. The extent of A30 in the mice identified in this study is consistent with that identified with molecular markers such as *Penk* (see figure 2 of Hu et al., 2020) and *Ddit4l, Npnt, Fam3c, Hlf, Epha4* and *Vamp1* (see www.brain-map.org). It is obvious that the rats and mice have a relatively huge A29 and much smaller A30 and A23~ (see Fig. 1). The connectivity of A29, A30 and A23~ in the rodents is different from one anther. For example, A29 (RSg) receives its inputs mainly from the AD, AV, LD, Cla, ACA (A24), A30 (RSag), PrS and Sub (Wyss and van Groen, 1992; Ding et al., 2020) and projects mainly to the AD, AV, LD, A24, A23~, PrS-PoS and PaS (Wyss and van Groen, 1992; Ding, 2013). A30 (RSag) gets its inputs mainly from the LD, AD, AV, M2, V2M, V1, V2L and PoR, (van Groen and Wyss, 1992; also see https://connectivity.brain-map.org) and innervates the LD, A29, visual (V2M, V2L and V1) and auditory (A2 and the posterior TeA) cortices as well as the PrS-PoS, PaS and PoR (Wyss and van Groen, 1992; Ding, 2013; the present study). A23~ in the rodents obtains its afferent inputs mainly from the AM, LP-Pul (medial part), A29, ACA and LD, and mainly targets the AM, LP-Pul, ACA, Orbm, PrS-PoS, SC (this study). Taken together, A29, A30 and A23~ have differential molecular signature and connectivity and thus likely play different roles in spatial processing, navigation, and adaptive behaviors.

## 4.4 Comparison of area 23 with parietal association cortex

In the primates, the PtA or area 7 (A7) mainly contains areas 7a (PG), 7b (PF), 7ip, 7m (PGm) and Opt (Leichnetz, 2001; Seltzer and Pandya, 2009). One of the major differences between PtA/A7 and A23 is the origins of thalamic inputs. As discussed above, A23 is strongly innervated by AM inputs. In contrast, PtA/A7 did not receive significant inputs from the AM (Schmahmann and Pandya, 1990; Leichnetz, 2001; Buckwalter et al., 2008). The second difference is about the projections to the RS. PtA/A7 appears to send strong projections to A30 while A23 has strong connections with A29 (Mesulam et al., 1977; Cavada and Goldman-Rakic, 1989; Kobayashi and Amaral, 2003). The third one is about its outputs to the PrS. PtA/A7 in general has stronger outputs than A23 does (Seltzer and Van Hoesen, 1979; Pandya et al., 1981; Ding et al., 2000). Additional difference is on the projections to the Cla with A23 and A7 displaying stronger (Van Hoesen et al., 1993; Parvizi et al., 2006) and weaker (Weber and Yin, 1984; Baizer et al., 1993) efferent projections to the Cla, respectively. All these differences also appear true for rodents. First, the PtA or posterior parietal cortex does not appear to receive the AM inputs in rats (Reep et al., 1994; Torrealba and Valdés, 2008; Olsen and Witter, 2016) while A23~ does (the present study). Second, the PtA and A23~ have stronger connections with RSag (A30) and RSg (A29), respectively (Wilber et al., 2015; the present study). Third, the PtA and A23~ send stronger and weaker projections to the PrS, respectively (Ding, 2013; Olsen et al., 2017; the present study). Fourth, the PtA and A23~ have weaker and stronger connections with Cla, respectively (Lyamzin and Benucci, 2019; the present study). It should be mentioned that the rodent PtA roughly contains the VISal, VISam, VISrl and the rostral part of the RSagl in Wang et al. (2020) (Carey and Neal, 1985; Reep et al., 1994; Swanson, 2004; Torrealba and Valdes, 2008; Paxinos and Watson, 2013; Olsen and Witter, 2016; Hovde et al., 2019; Lyamzin and Benucci, 2019; Gilissen et al., 2021).

## 4.5 Functional consideration of area 23

Since A23 was not reported in the rodents, previous functional studies of A23 or PCC were mainly conducted in human and NHP brains. Briefly, A23 is a key component of the DMN and displays increased activity when retrieving episodic and autobiographical memories, planning for future, understanding the behavior and thoughts of others, appraising emotional information and “free-wheeling” during unconstrained “rest” (Leech and Sharp, 2014; Rolls, 2019; Zhang et al., 2019). In addition, A23 is sensitive to the changes in arousal state, awareness and the environment and participates in controlling the balance between internal and external attention (Leech and Sharp, 2014; Barack and Platt, 2021; Chang et al., 2021). Connectional studies in the NHP appear to support these functions since A23 converges many aspects of the inputs from hetero-modal cortical and subcortical regions.

For instance, the monkey A23 receives direct inputs from multimodal association cortices such as visual (areas 18, 19 and TPO), parietal (areas 7a and Opt), auditory (areas TB, TA and Tpt) and parahippocampal (areas TH and TF) association cortices. A23 is also innervated by different limbic cortices such as the ORBm (areas 11, 13 and 14), ACC (A24 and A32), subcallosal cingulate cortex (area 25), RS (A29 and A30), PRC (areas 35 and 36) and entorhinal cortex (MEC and LEC) (Baleydier and Mauguiere, 1980; Vogt and Pandya, 1987; Van Hosen et al., 1993; Yukie, 1995; Parvizi et al., 2006; Seltzer and Pandya, 2009; Balcerek et al., 2021). In addition, the DFC (areas 8, 9, 10, 46) also sends projections to A23 (Vogt and Pandya, 1987; Kobayashi and Amaral, 2003; Parvizi et al., 2006). Most of these connections have been confirmed in the rodent A23~ in the present study except for the DFC, which remains to be explored in the rodents. Therefore, A23 has access to a variety of information from different modalities and thus is a critical hub for integration and regulation of different information.

Other important information also reaches A23. For example, the ORBm in both monkeys and rodents displays strong reciprocal connections with A23 (Vogt and Pandya, 1987; Van Hosen et al., 1993; Parvizi et al., 2006; the present study). Specialized areas for value updating (area 13) and goal selections (area 11) were reported in monkey ORBm (e.g., Murray et al., 2015) and this may be also true for human ORBm (Saez et al., 2018). The ORBm may be a critical site for temporal cognition, integrating reward magnitude and delays, which are needed for value processing (Sosa et al., 2021). The value-related information in the ORBm would project to and interact with A23 and thus could contribute to the functions of A23 in planning for future and understanding the behavior and thoughts of others.

The RS (A29 and A30) is another region with heavy connection with A23 in both monkeys (Morris et al., 1999; Kobayashi and Amaral, 2003, 2007) and rodents (the present study). For instance, A29 in the rodents receive its major inputs from the Sub, PrS-PoS and A30 as well as the AD, AV and LD, all of which are heavily involved in spatial processing and navigation (Wyss and van Groen, 1992; sugar et al., 2011; Ding et al., 2020; the present study). A29 mainly projects to the PrS-PoS, A30, PaS, ACA (A24), AD, AV and LD (Wyss and van Groen, 1992) and A23 (the present study). In contrast, A30 appears to converge sensory, motor and spatial information since A30 receives dense inputs from the primary (V1) and secondary (V2M and V2L) visual inputs (Wyss and van Groen, 1992; the present study) as well as from the secondary motor area M2 (the present study), LD, LP-Pul and PtA (Torrealba and Valdés, 2008). A30 projects back to these afferent regions and to the PrS-PoS and A29 (the present study). Although A30 in the rodents does not appear to have strong connections with A23, it heavily projects to A29, which strongly innervate A23. Therefore, A23 could access different spatial information about the environment via the RS and enable its roles in spatial learning and memory such as retrieving episodic and autobiographical memories and adapting changes in the environment. Other aspects of memory-related information could reach A23 via the AM, PoR, LEC and MEC, all of which receive dense projections from the Sub and/or ProS. The latter two are the output structures of the hippocampal memory system (see Ding et al., 2020).

The ACC is heavily and reciprocally connected with A23, RS, ORBm, PoR, AM, BL and Cla (Baleydier and Mauguiere, 1980; Vogt et al., 1987; Vogt and Pandya, 1987; Van Hosen et al., 1993; Kobayashi and Amaral, 2003). The ACC probably carries a myriad of signals such as error detection, reinforcement/feedback, value, response conflict, autonomic and emotional states, which are necessary for the modulation of attention and task-relevant/irrelevant signals so that difficult decisions can be made and action plans adapted when necessary (Rolls, 2019; Brockett and Roesch, 2021; Seamans and Floresco, 2022). All these information can reach to and interact with A23 via the strong reciprocal connections between A23 and the anterior cingulate cortex (for monkeys: Baleydier and Mauguiere, 1980; Van Hosen et al., 1993; Parvizi et al., 2006; Rolls, 2019; for rodents: the present study).

Finally, A23 in both primates and rodents is also reciprocally and heavily connected with the Cla (Baleydier and Mauguiere, 1980; Van Hosen et al., 1993; Parvizi et al., 2006; the present study). Many previous studies reported that the Cla is heavily involved in attention, awake-sleep states, sensory and salience processing likely via regulation of cortical excitability (see review in Smith et al., 2020). Recent works indicate that Cla mainly exerts inhibitory roles on widespread cortical regions including cingulate cortex (Smith et al., 2020). Therefore, as an example, it is likely that the Cla could inhibit or interacts with the DMN (including A24, A23 and others) when salient stimuli appear in the environment so that the focus of attention can be switched to more salient tasks.

In summary, the PCC (A23) is a critical hub for the integration and modulation of multimodal information underlying spatial processing, self-reflection, attention, plan for future and many adaptive behaviors.

## Abbreviations

3V: third ventricle
4V: fourth ventricle
A29: area 29
A29l: lateral division of area 29
A29m: medial division of area 29
A30: area 30
A23: area 23
A23~: rodent equivalent of area 23
ac: anterior commissure
ACA: anterior cingulate area
AD: anterodorsal nucleus of thalamus
AM: anteromedial nucleus of thalamus
Aq: aqueduct
AuD: auditory cortex
AV: anteroventral nucleus of thalamus
BL: basolateral amygdaloid nucleus
cc: corpus callosum
ccs: splenium of corpus callosum
Cla: claustrum
CPu: caudate putamen
CPu-p: posterior part of caudate putamen
DFC: dorsolateral prefrontal cortex
DLG: dorsal lateral geniculate nucleus
EC: entorhinal cortex
fr: fasciculus retroflexus
Hip: hippocampal formation
IG: indusium griseum
IL: intralimbic cortex
LD: lateral dorsal nucleus of thalamus
LEC: lateral entorhinal cortex
LP-Pul: lateroposterior nucleus-pulvinar complex
LS: lateral septal nucleus
LV: lateral ventricle
M2: secondary motor cortex
MEC: medial entorhinal cortex
NAC: nucleus accumbens
NC: nucleus coeruleus
NLL: nucleus of lateral lemniscus
ORBm: medial orbitofrontal cortex
PAG: periaqueductal gray
PaS: parasubiculum
PHC: posterior parahippocampal cortex
PHN: posterior hypothalamic nucleus
PL: prelimbic cortex
PN: pontine nucleus
PoR: postrhinal cortex
PRC: perirhinal cortex
Pro: area prostriata
PrS: presubiculum
PrSd: dorsal presubiculum (postsubiculum)
PrSv: ventral presubiculum
PT: paratenial nucleus
PtA: parietal association cortex
PTN: pretectal nucleus
PV: paraventricular hypothalamic nucleus
PVT: paraventricular nucleus of thalamus
Re: reuniens nucleus
RN: red nucleus
RS: retrosplenial cortex
RSag: agranular retrosplenial cortex
RSagl: lateral agranular retrosplenial cortex
RSg: granular retrosplenial cortex
RT: reticular thalamic nucleus
SC: superior colliculus
Sd: dorsal subiculum
smt: stria medullaris of thalamus
Spo: polymorphic layer of subiculum
Sub: subiculum
SuS: supracallosal subiculum
TeA: temporal association cortex
V1: primary visual cortex
V2L: lateral secondary visual cortex
V2M: medial secondary visual cortex
VA: ventroanterior nucleus
VLG: ventral lateral geniculate nucleus
VLG-m: medial part of VLG
wm: white matter
ZI: zona incerta
ZIp: posterior part of zona incerta
ZIa: anterior part of zona incerta

## Data availability statement

The original data from the rats are included in this article. Raw data on the mouse connectivity are publicly available (https://connectivity.brain-map.org). Further inquiries can be directed to the corresponding author.

## Ethics statement

This animal study was reviewed and approved by Institutional Animal Care and Use Committee of the Guangzhou Medical University.

## Author contributions

SLD: conceptualization. XJX, SLD, SQC, CHC and SYZ: investigation. XJX and SLD: manuscript writing. SLD, SQC and XQZ: supervision. All authors read and approved the final manuscript.

## REFERENCES

Baizer, J. S., Desimone, R., and Ungerleider, L. G. (1993) Comparison of subcortical connections of inferior temporal and posterior parietal cortex in monkeys. Vis Neurosci 10, 59–72.

Balcerek, E., Włodkowska, U., and Czajkowski, R. (2021) Retrosplenial cortex in spatial memory: focus on immediate early genes mapping. Mol Brain 14, 172. doi: 10.1186/s13041-021-00880-w.

Baleydier, C., and Mauguiere, F. (1980) The duality of the cingulate gyrus in monkey. Neuroanatomical study and functional hypothesis. Brain 103, 525–554.

Baleydier, C., and Mauguiere, F. (1985) Anatomical evidence for medial pulvinar connections with the posterior cingulate cortex, the retrosplenial area, and the posterior parahippocampal gyrus in monkeys. J Comp Neurol 232, 219–228.

Barack, D.L., and Platt, M.L. (2021) Neuronal Activity in the Posterior Cingulate Cortex Signals Environmental Information and Predicts Behavioral Variability during Trapline Foraging. J Neurosci 41, 2703–2712.

Bluhm, R. L., Miller, J., Lanius, R. A., Osuch, E. A., Boksman, K., Neufeld, R. W. J., Théberge, J., Schaefer, B., and Williamson, P. C. (2009) Retrosplenial cortex connectivity in schizophrenia. Psychiatry Res 174, 17–23.

Brodmann, K. (1909) Vergleichende lokalisationslehre der Grosshirnrinde. Barth.

Brockett, A. T., and Roesch, M. R. (2021) Anterior cingulate cortex and adaptive control of brain and behavior. Int Rev Neurobiol 158, 283–309.

Buckner, R. L., Andrews-Hanna, J. R., and Schacter, D. L. (2008) The brain’s default network: anatomy, function, and relevance to disease. Ann N Y Acad Sci 1124, 1–38.

Buckwalter, J. A., Parvizi, J., Morecraft, R. J., and van Hoesen, G. W. (2008) Thalamic projections to the posteromedial cortex in the macaque, J Comp Neurol 507, 1709–1733.

Carey, R.G., and Neal, T.L., (1985) The rat claustrum: afferent and efferent connections with visual cortex. Brain Res 329, 185–193.

Cavada, C., and Goldman-Rakic, P.S. (1989) Posterior parietal cortex in rhesus monkey: I. Parcellation of areas based on distinctive limbic and sensory corticocortical connections. J Comp Neurol 287, 393–421.

Chang, D.H.F., Jiang, B., Wong, N.H.L., Wong, J.J., Webster, C., Lee, T.M.C. (2021) The human posterior cingulate and the stress-response benefits of viewing green urban landscapes. Neuroimage 226, 17555. doi: 10.1016/j.neuroimage.2020.117555.

Chen, C.-H., Hu, J.-M., Zhang, S.-Y., Xiang, X.-J., Chen, S.-Q., and Ding, S.-L. (2021) Rodent area prostriata converges multimodal hierarchical inputs and projects to the structures important for visuomotor behaviors. Front Neurosci 15, 772016.

Chen, S.Q., Chen, C.H., Xiang, X.J., Zhang, S.Y., and Ding, S.L. (2022) Chemoarchitecture of area prostriata in adult and developing mice: comparison with presubiculum and parasubiculum. J Comp Neurol 530, 2486–2517. doi: 10.1002/cne.25346.

de Lima, M. A. X., Baldo, M. V. C., and Canteras, N. S. (2019) Revealing a Cortical Circuit Responsive to Predatory Threats and Mediating Contextual Fear Memory. Cereb Cortex 29, 3074–3090.

Dean, H. L., Crowley, J. C., and Platt, M. L. (2004) Visual and saccade-related activity in macaque posterior cingulate cortex. J Neurophysiol 92, 3056–3068.

Dierssen, G., Odoriz, B., and Hernando, C. (1969) Sensory and motor responses to stimulation of the posterior cingulate cortex in man. J Neurosurg 31, 435–440.

Dillen, K. N. H., Jacobs, H. I. L., Kukolja, J., von Reutern, B., Richter, N., Onur, Ö. A., Dronse, J., Langen, K.-J., and Fink, G. R. (2016) Aberrant functional connectivity differentiates retrosplenial cortex from posterior cingulate cortex in prodromal Alzheimer’s disease. Neurobiol Aging 44, 114–126.

Ding, S.-L. (2013) Comparative anatomy of the prosubiculum, subiculum, presubiculum, postsubiculum, and parasubiculum in human, monkey, and rodent. J Comp Neurol 521, 4145–4162.

Ding, S.L. (2022) A novel subdivision of the bed nucleus of stria terminalis in monkey, rat and mouse brains. J Comp Neurol, 1–25. https://doi.org/10.1002/cne.25446.

Ding, S. L., Van Hoesen, G., and Rockland, K. S. (2000) Inferior parietal lobule projections to the presubiculum and neighboring ventromedial temporal cortical areas. J Comp Neurol 425, 510–530.

Ding, S. L., and Van Hoesen, G. W. (2010) Borders, extent, and topography of human perirhinal cortex as revealed using multiple modern neuroanatomical and pathological markers. Hum Brain Mapp 31, 1359–1379. doi: 10.1002/hbm.20940.

Ding, S. L., and Van Hoesen, G. W. (2015) Organization and detailed parcellation of human hippocampal head and body regions based on a combined analysis of cyto-and chemoarchitecture. J Comp Neurol 523, 2233–2253. doi: 10.1002/cne.23786.

Ding, S.-L., Royall, J. J., Sunkin, S. M., Ng, L., Facer, B. A. C., Lesnar, P., Guillozet-Bongaarts, A., McMurray, B., Szafer, A., Dolbeare, T. A., Stevens, A., Tirrell, L., Benner, T., Caldejon, S., Dalley, R. A., Dee, N., Lau, C., Nyhus, J., Reding, M., Riley, Z. L., Sandman, D., Shen, E., van der Kouwe, A., Varjabedian, A., Wright, M., Zöllei, L., Dang, C., Knowles, J. A., Koch, C., Phillips, J. W., Sestan, N., Wohnoutka, P., Zielke, H. R., Hohmann, J. G., Jones, A. R., Bernard, A., Hawrylycz, M. J., Hof, P. R., Fischl, B., and Lein, E. S. (2016) Comprehensive cellular-resolution atlas of the adult human brain. J Comp Neurol 524, 3127–3481.

Ding, S.-L., Yao, Z., Hirokawa, K. E., Nguyen, T. N., Graybuck, L. T., Fong, O., Bohn, P., Ngo, K., Smith, K. A., Koch, C., Phillips, J. W., Lein, E. S., Harris, J. A., Tasic, B., and Zeng, H. (2020) Distinct transcriptomic cell types and neural circuits of the subiculum and prosubiculum along the dorsal-ventral axis. Cell Rep 31, 107648.

Ding, S.L., Royall, J.J., Lesnar, P., Facer, B.A.C., Smith, K.A., Wei, Y., Brouner, K., Dalley, R.A., Dee, N., Dolbeare, T.A., Ebbert, A., Glass, I.A., Keller, N.H., Lee, F., Lemon, T.A., Nyhus, J., Pendergraft, J., Reid, R., Sarreal, M., Shapovalova, N.V., Szafer, A., Phillips, J.W., Sunkin, S.M., Hohmann, J.G., Jones, A.R., Hawrylycz, M.J., Hof, P. R., Ng, L., Bernard, A., Lein, E.S. (2022) Cellular resolution anatomical and molecular atlases for prenatal human brains. J Comp Neurol 530, 6–503. doi: 10.1002/cne.25243.

Fudge, J. L., Breitbart, M. A., and McClain, C. (2004) Amygdaloid inputs define a caudal component of the ventral striatum in primates. J Comp Neurol 476, 330–347.

Gehrlach, D. A., Weiand, C., Gaitanos, T. N., Cho, E., Klein, A. S., Hennrich, A. A., Conzelmann, K.-K., and Gogolla, N. (2020) A whole-brain connectivity map of mouse insular cortex. Elife 9, e55585. doi: 10.7554/eLife.55585.

Gilissen, S. R. J., Farrow, K., Bonin, V., and Arckens, L. (2021) Reconsidering the border between the visual and posterior parietal cortex of mice. Cereb Cortex 31, 1675–1692.

Gonzalo-Ruiz, A., Sanz, J. M., Morte, L., and Lieberman, A. R. (1997) Glutamate and aspartate immunoreactivity in the reciprocal projections between the anterior thalamic nuclei and the retrosplenial granular cortex in the rat. Brain Res Bull 42, 309–321.

Hovde, K., Gianatti, M., Witter, M. P., and Whitlock, J. R. (2019) Architecture and organization of mouse posterior parietal cortex relative to extrastriate areas. Eur J Neurosci 49, 1313–1329.

Hsu, L.-M., Liang, X., Gu, H., Brynildsen, J. K., Stark, J. A., Ash, J. A., Lin, C.-P., Lu, H., Rapp, P. R., Stein, E. A., and Yang, Y. (2016) Constituents and functional implications of the rat default mode network. Proc Natl Acad Sci U S A 113, E4541–E4547.

Hu, J.-M., Chen, C.-H., Chen, S.-Q., and Ding, S.-L. (2020) Afferent Projections to Area Prostriata of the Mouse. Front Neuroanat 14, 605021.

Kobayashi, Y., and Amaral, D. G. 2003. Macaque monkey retrosplenial cortex: III. Cortical afferents. J Comp Neurol 466, 48–79.

Kobayashi, Y., and Amaral, D. G. 2007. Macaque monkey retrosplenial cortex: III. Cortical efferents. J Comp Neurol 502, 810–833.

Leech, R. and Sharp, D.J. (2014) The role of the posterior cingulate cortex in cognition and disease. Brain 137, 12–32. doi: 10.1093/brain/awt162.

Leichnetz, G. R. (2001) Connections of the medial posterior parietal cortex (area 7m) in the monkey. Anat Rec 263, 215–236.

Leung, M.-K., and Lau, W. K.-W. (2020) Resting-state abnormalities of posterior cingulate in autism spectrum disorder. Prog Mol Biol Transl Sci 173, 139–159.

Lu, H., Zou, Q., Gu, H., Raichle, M. E., Stein, E. A., and Yang, Y. (2012) Rat brains also have a default mode network. Proc Natl Acad Sci U S A 109, 3979–3984.

Lu, W., Chen, S., Chen, X., Hu, J., Xuan, A., and Ding, S.L. (2020). Localization of area prostriata and its connections with primary visual cortex in rodent. J Comp Neurol. 528, 389-406. doi: 10.1002/cne.24760.

Lyamzin, D., and Benucci, A. (2019) The mouse posterior parietal cortex: Anatomy and functions. Neurosci Res 140, 14–22.

Mantini, D., Gerits, A., Nelissen, K., Durand, J.-B., Joly, O., Simone, L., Sawamura, H., Wardak, C., Orban, G. A., Buckner, R. L., and Vanduffel, W. (2011) Default mode of brain function in monkeys. J Neurosci 31, 12954–12962.

Mesulam, M.M.,Van Hoesen, G.W., Pandya, D.N., and Geschwind, N. (1977) Limbic and sensory connections of the inferior parietal lobule (area PG) in the rhesus monkey: a study with a new method for horseradish peroxidase histochemistry. Brain Res 136, 393–414.

Miller, M.W., and Vogt, B.A. (1984) Direct connections of rat visual cortex with sensory, motor, and association cortices. J Comp Neurol 226, 184–202. doi: 10.1002/cne.902260204.

Morecraft, R. J., Cipolloni, P. B., Stilwell-Morecraft, K. S., Gedney, M. T., and Pandya, D. N. (2004) Cytoarchitecture and cortical connections of the posterior cingulate and adjacent somatosensory fields in the rhesus monkey. J Comp Neurol 469, 37–69.

Morris, R., Petrides, M., and Pandya, D. N. (1999) Architecture and connections of retrosplenial area 30 in the rhesus monkey (Macaca mulatta). Eur J Neurosci 11, 2506–2518.

Murray, E. A.,Moylan, E.J., Saleem, K.S., Basile, B.M., and Turchi, J. (2015) Specialized areas for value updating and goal selection in the primate orbitofrontal cortex. Elife 4, e11695.

Olsen, G. M., Ohara, S., Iijima, T., and Witter, M. P. (2017) Parahippocampal and retrosplenial connections of rat posterior parietal cortex. Hippocampus 27, 335–358.

Olsen, G. M., and Witter, M. P. (2016) Posterior parietal cortex of the rat: Architectural delineation and thalamic differentiation. J Comp Neurol 524, 3774–3809.

Pandya, D. N., Van Hoesen, G. W., and Mesulam, M. M. (1981) Efferent connections of the cingulate gyrus in the rhesus monkey. Exp Brain Res 42, 319–330.

Parvizi, J., Van Hoesen, G.W., Buckwalter, J., and Damasio, A. (2006) Neural connections of the posteromedial cortex in the macaque. Proc Natl Acad Sci U S A 103, 1563–1568. doi: 10.1073/pnas.0507729103.

Paxinos, G., and Watson, C. (2013) The Rat Brain in Stereotaxic Coordinates. San Diego, CA: Academic Press.

Paxinos, G., & Franklin, K.B.J. (2012) The mouse Brain in Stereotaxic Coordinates. San Diego, CA: Academic Press.

Raichle, M. E. (2015) The brain’s default mode network. Annu Rev Neurosci 38, 433–447.

Reep, R. L., Chandler, H. C., King, V., and Corwin, J. V. (1994) Rat posterior parietal cortex: topography of corticocortical and thalamic connections. Exp Brain Res 100, 67–84.

Rolls, E. T. (2019) The cingulate cortex and limbic systems for action, emotion, and memory. Handb Clin Neurol 166, 23–37.

Rolls, E. T., and Wirth, S. (2018) Spatial representations in the primate hippocampus, and their functions in memory and navigation. Prog Neurobiol 171, 90–113.

Saez, I., Lin, J., Stolk, A., Chang, E., Parvizi, J., Schalk, G., Knight, R. T., and Hsu, M. (2018) Encoding of Multiple Reward-Related Computations in Transient and Sustained High-Frequency Activity in Human OFC. Curr Biol 28, 2889–2899.

Schmahmann, J. D., and Pandya, D. N. (1990) Anatomical investigation of projections from thalamus to posterior parietal cortex in the rhesus monkey: a WGA-HRP and fluorescent tracer study. J Comp Neurol 295, 299–326.

Seamans, J. K., and Floresco, S. B. (2022) Event-based control of autonomic and emotional states by the anterior cingulate cortex. Neurosci Biobehav Rev 133, 104503. doi: 10.1016/j.neubiorev.2021.12.026.

Seltzer, B., and Van Hoesen, G. W. (1979) A direct inferior parietal lobule projection to the presubiculum in the rhesus monkey. Brain Res 179, 157–161.

Seltzer, B., and Pandya, D. N. (2009) Posterior cingulate and retrosplenial cortex connections of the caudal superior temporal region in the rhesus monkey. Exp Brain Res 195, 325–334.

Shibata, H. (1993) Efferent projections from the anterior thalamic nuclei to the cingulate cortex in the rat. J Comp Neurol 330, 533–542.

Shibata, H., and Yukie, M. (2003) Differential thalamic connections of the posteroventral and dorsal posterior cingulate gyrus in the monkey. Eur J Neurosci 18, 1615–1626.

Smith, J.B., Lee, A.K., and Jackson, J. (2020) The Claustrum. Cur Biol 30, R1391–R1412.

Sosa, J.L.R., Buonomano, D., and Izquierdo, A. (2021) The orbitofrontal cortex in temporal cognition. Behav Neurosci 135, 154–164.

Sripanidkulchai, K., and Wyss, J. M. (1986) Thalamic projections to retrosplenial cortex in the rat. J Comp Neurol 254, 143–165.

Sugar, J., Witter, M. P., van Strien, N. M., and Cappaert, N. L. M. (2011) The retrosplenial cortex: intrinsic connectivity and connections with the (para)hippocampal region in the rat. An interactive connectome, Front Neuroinform 5, 7.

Swanson, L.W. (2004) Brain maps: structure of the rat brain (third edition). Amsterdam, The Netherlands: Elsevier.

Torrealba, F., and Valdés, J. L. (2008) The parietal association cortex of the rat. Biol Res 41, 369–377.

van Groen, T., Kadish, I., and Wyss, J. M. (1999) Efferent connections of the anteromedial nucleus of the thalamus of the rat. Brain Res Brain Res Rev 30, 1–26.

van Groen, T., and Wyss, J. M. (1992) Connections of the retrosplenial dysgranular cortex in the rat, J Comp Neurol 315, 200–216.

van Hoesen, G.W., Morecraft, R.J., and Vogt, B.A. (1993) Connections of the monkey cingulate cortex. In book: Neurobiology of Cingulate Cortex and Limbic Thalamus (Editors: Vogt BA and Gabriel M). publisher, Birkhauser, Boston, MA. pp249–284.

Vertes, R.P., and Hoover, W.B. (2008) Projections of the paraventricular and paratenial nuclei of the dorsal midline thalamus in the rat. J Comp Neurol 508, 212–237.

Vertes, R.P., Hoover, W.B., Do Valle, A.C., Sherman, A., and Rodriguez, J.J. (2006) Efferent projections of reuniens and rhomboid nuclei of the thalamus in the rat. J Comp Neurol 499, 768–796.

Vogt, B. A., and Miller, M. W. (1983) Cortical connections between rat cingulate cortex and visual, motor, and postsubicular cortices, J Comp Neurol 216, 192–210.

Vogt, B. A., and Pandya, D. N. (1987) Cingulate cortex of the rhesus monkey: II. Cortical afferents, J Comp Neurol 262, 271–289.

Vogt, B. A., Pandya, D. N., and Rosene, D. L. (1987) Cingulate cortex of the rhesus monkey: I. Cytoarchitecture and thalamic afferents, J Comp Neurol 262, 256–270.

Vogt, B.A., Vogt, L., Perl, D.P., Hof, P. R. (2001) Cytology of human caudomedial cingulate, retrosplenial, and caudal parahippocampal cortices. J Comp Neurol 438, 353–376. doi: doi: 10.1002/cne.1320.

Vogt, B.A., Vogt, L., Farber, N.B., and Bush, G. (2005) Architecture and neurocytology of monkey cingulate gyrus. J Comp Neurol 485, 218–239. doi: 10.1002/cne.20512.

Vogt, B.A., and Paxinos, G. (2014) Cytoarchitecture of mouse and rat cingulate cortex with human homologies. Brain Struct Funct 219, 185–192.

Wallén-Mackenzie, Å., Dumas, S., Papathanou, M., Martis Thiele, M.M., Vlcek, B., König, N., Björklund, Å.K. (2020) Spatio-molecular domains identified in the mouse subthalamic nucleus and neighboring glutamatergic and GABAergic brain structures. Commun Biol 3, 338. doi: 10.1038/s42003-020-1028-8.

Wang, Q., Ding, S.-L., Li, Y., Royall, J., Feng, D., Lesnar, P., Graddis, N., Naeemi, M., Facer, B., Ho, A., Dolbeare, T., Blanchard, B., Dee, N., Wakeman, W., Hirokawa, K. E., Szafer, A., Sunkin, S. M., Oh, S. W., Bernard, A., Phillips, J. W., Hawrylycz, M., Koch, C., Zeng, H., Harris, J. A., and Ng, L. (2020) The Allen Mouse Brain Common Coordinate Framework: A 3D Reference Atlas, Cell 181, 936–953.

Weber, J.T., and Yin, T.C. (1984) Subcortical projections of the inferior parietal cortex (area 7) in the stump-tailed monkey. J Comp Neurol 224, 206–230.

Wilber, A.A., Clark, B.J., Demecha, A.J., Mesina, L., Vos, J.M., McNaughton, B.L. (2015) Cortical connectivity maps reveal anatomically distinct areas in the parietal cortex of the rat. Front Neural Circuits 8, 146. doi: 10.3389/fncir.2014.00146.eCollection2014.

Wyss, J.M., and van Groen, T. (1992) Connections between the retrosplenial cortex and the hippocampal formation in the rat: a review. Hippocampus 2, 1–11. doi: 10.1002/hipo.450020102.

Yamawaki, N., Radulovicm, J., Gordon, M.G., and Shepherd, G.M.G. (2016) A corticocortical circuit directly links retrosplenial cortex to M2 in the mouse. J Neurosci 36, 9365–9374.

Yukie, M. (1995). Neural connections of auditory association cortex with the posterior cingulate cortex in the monkey. Neurosci Res 22,179–187.

Zhang, R., and Volkow, N.D. (2019) Brain default-mode network dysfunction in addiction. NeuroImage 200, 313–331.

